# Distinct dopamine circuits transmit the reinforcing and anxiogenic effects of nicotine

**DOI:** 10.1101/2020.07.30.228189

**Authors:** C Nguyen, S Mondoloni, I Centeno, R Durand-de Cuttoli, S Tolu, S Valverde, T Le Borgne, B Hannesse, S Pons, U Maskos, D Dalkara, JP Hardelin, A Mourot, F Marti, P Faure

## Abstract

Nicotine, the addictive component of tobacco, stimulates dopamine (DA) neurons of the ventral tegmental area (VTA) to establish and maintain reinforcement. Nicotine also induces negative emotional states such as anxiety, yet through an unknown circuitry. Here we show that nicotine at reinforcing doses drives opposite functional responses on two distinct populations of VTA DA neurons with anatomically segregated projections: it activates those that project to the nucleus accumbens (NAc) while it inhibits those that project to the amygdala nuclei (Amg). We further show that nicotine, by acting on β2 subunit-containing nicotinic acetylcholine receptors of the VTA, mediates both reinforcement and anxiety. Finally, using optogenetic experiments we dissociate the roles of the VTA-NAc excitation and VTA-Amg inhibition in reinforcement and anxiety-like behavior, respectively. We thus propose that the positive and negative behavioral outcomes of nicotine consumption involve distinct subpopulations of VTA DA neurons with opposite responses to nicotine.

## Introduction

Nicotine is the principal addictive component that drives continued tobacco use. The initiation of addiction involves the mesocorticolimbic dopamine (DA) system, which contributes to the processing of rewarding stimuli during the overall shaping of successful behaviors (Schultz, 2007). It has been hypothesized that addictive drugs such as nicotine hijack the same mechanisms as reinforcement learning, leading to an overvaluation of the drug reward at the expense of natural rewards. While drug-induced reinforcement learning generally involves an increase in extracellular DA concentration in the nucleus accumbens (NAc), the underlying molecular and cellular mechanisms are drug dependent (Changeux, 2010; Di Chiara and Imperato, 1988; Luscher, 2016). Nicotine, for instance, exerts its reinforcing effects through the direct activation of nicotinic acetylcholine receptors (nAChRs), a family of pentameric ligand-gated ion channels (Changeux et al., 1998), on midbrain DA and GABA neurons, thus increasing the activity of both neuronal populations (Maskos et al., 2005; Morel et al., 2014; Tolu et al., 2013). Cell type-specific optogenetic manipulations have confirmed that DA cell activation is sufficient to drive the transition toward addiction, and established causal links between DA neuron activation and drug-adaptive behaviors (Pascoli et al., 2015). However, such view does not take into account the heterogeneity of midbrain DA neurons and the possibility that different messages can be transmitted in parallel from DA neurons of the ventral tegmental area (VTA). Indeed, VTA DA neurons belong to anatomically distinct circuits, differ in their molecular properties, and show diverse responses to external stimuli (Lammel et al., 2008; Poulin et al., 2018). DA neurons not only transmit signals related to salience and reward, but also to aversive stimuli (Brischoux et al., 2009; de Jong et al., 2019), including the negative effects of nicotine at high doses (Grieder et al., 2019; 2010). However, how DA neurons simultaneously transmit opposing signals in response to the same stimuli remains unclear. Whereas the vast majority of research teams that have examined nicotine-evoked responses report a homogenous activation of DA neurons and an increase in DA release in their projection areas (Di Chiara and Imperato, 1988; Grenhoff et al., 1986; Mansvelder and McGehee, 2000; Maskos et al., 2005; Picciotto et al., 1998; Zhao-Shea et al., 2011), other reports suggested that DA neuron responses to nicotine are more heterogeneous than previously thought (Eddine et al., 2015; Mameli-Engvall et al., 2006; Zhao-Shea et al., 2011). Therefore, a key issue is how the multiple effects of nicotine map onto the DA cell diversity, and whether nAChRs or other features can define different neuronal subpopulations that, through their response to nicotine, would influence specific behaviors.

## Results

### DA neurons in the VTA display opposite responses to acute nicotine injection

We performed single-cell electrophysiological recordings on anaesthetized mice, using an intravenous (IV) injection of a reinforcing dose of nicotine (30 μg/kg) (Morel et al., 2014), to record the responses of VTA DA neurons (n = 245). These neurons were first identified during the recording based on their electrophysiological properties (i.e., firing rate and action potential width (Mameli-Engvall et al., 2006; Ungless and Grace, 2012)), and then filled with neurobiotin by the juxtacellular labeling technique (Eddine et al., 2015; Pinault, 1996). All neurons were confirmed as DA neurons *post-hoc* by immunofluorescence co-labeling with tyrosine hydroxylase (TH) (Figure 1A). Acute nicotine injections induced a significant variation, either positive or negative, of DA neuron firing rates (Figure 1B, left). Among the 245 DA neurons identified, some were activated (n = 155) whereas others were inhibited (n = 90) by the nicotine injection compared to the control saline injection, as previously reported (Eddine et al., 2015). The amplitudes of nicotine-induced activation and inhibition were similar, at about ± 35 % from baseline (Figure 1B, right). We first sought to determine whether the opposing responses to nicotine injection between these DA neuron populations were associated with a difference in their spontaneous activity. The spontaneous activity of VTA DA neurons is characterized by their firing rate and the percentage of spikes within a burst (%SWB) (Mameli-Engvall et al., 2006). Bursts are classically identified as discrete events consisting of a sequence of spikes with (i) burst onset defined by two consecutive spikes within an interval < 80 ms and (ii) burst termination defined by an inter-spike interval > 160 ms (Grace and Bunney, 1984a; Ungless and Grace, 2012). We found that DA neurons activated or inhibited by the nicotine injection had similar firing rates (Δ = 0.3 Hz, p = 0.054), however the inhibited neurons showed more bursting activity than the activated ones (Δ = 5.1 %, p = 0.04) (Figure 1C, left). An analysis of the distribution of burst time intervals also highlighted different profiles in the distribution of interspike intervals depending on the burst length (Figure 1C, right). Other parameters describing cell spontaneous activity (e.g. coefficient of variation, burst length, …) were analyzed, but none of them revealed a difference between activated and inhibited DA neurons (not shown). We thus observed small differences in bursting activity between activated and inhibited DA neurons, but we were not able to predict their nicotine-evoked responses based on the analysis of their spontaneous activity, either by traditional classification procedures or by machine learning (Figure S1 A - C). We next asked whether these two populations are anatomically segregated. Neurobiotin-filled cell bodies of each recorded neuron were positioned onto slices of the Mouse Brain Atlas (Paxinos and Franklin, 2004) (Figure S1 D) to study their anatomical location. As illustrated by a single atlas plate schematic (bregma - 3.3 mm), anatomical coordinates suggested that the inhibited neurons were located more medially within the VTA than the activated neurons, independently of their antero-posterior or dorso-ventral positions (Figure 1D).

**Figure 1:**
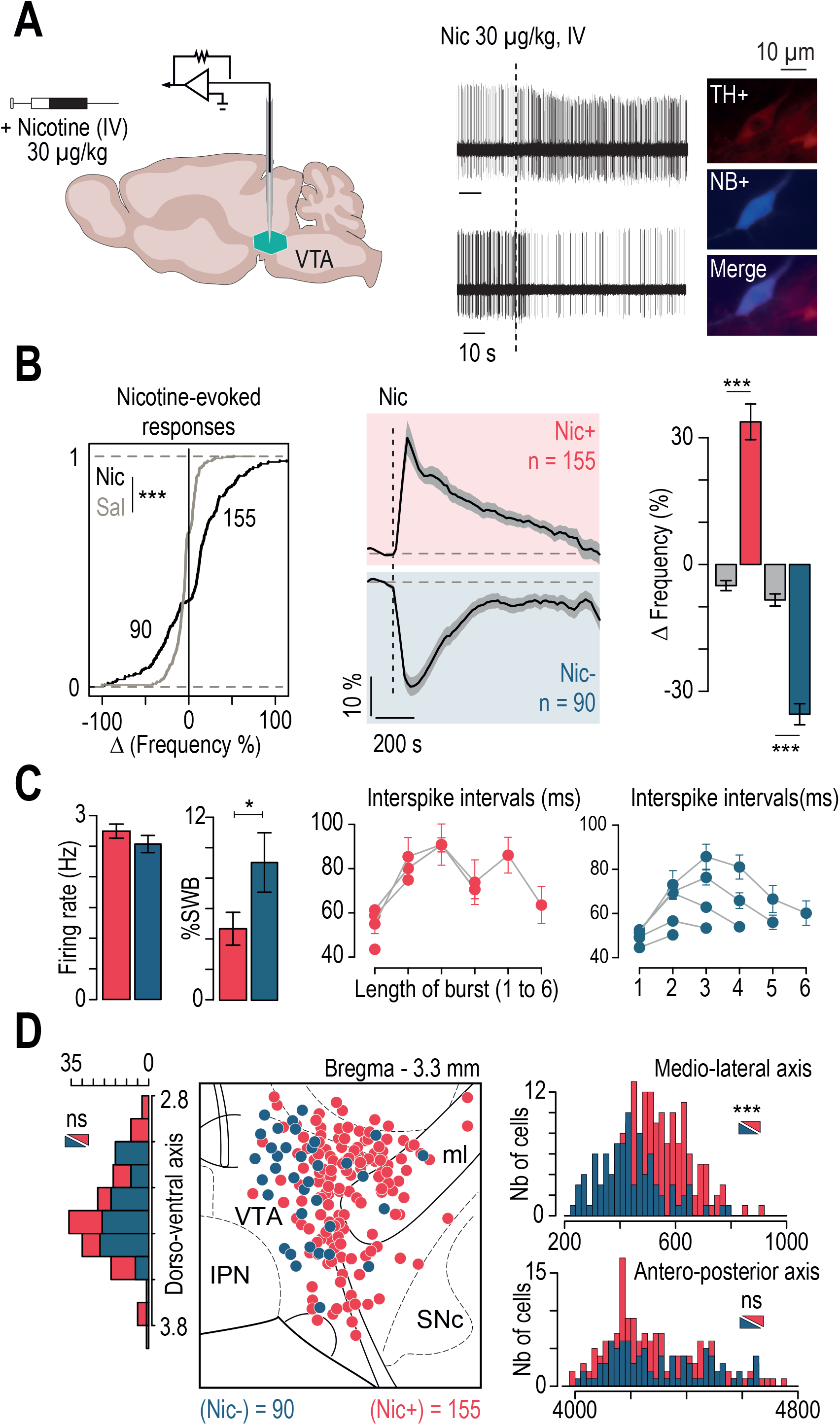
Nicotine evoked opposite responses in DA neurons of the VTA. **(A)** Intravenous (IV) injections of nicotine (Nic, 30 μg/kg) induce activation and inhibition of distinct VTA DA cells. Post-recording identification of neurobiotin (NB)-labeled VTA DA neurons by immunofluorescence (TH = tyrosine hydroxylase, NB = streptavidin-AMCA against neurobiotin). **(B)** *Left*: Cumulative distribution of responses after IV injection of either saline (Sal, grey, n = 233) or nicotine (Nic, black, n = 245) (Kolmogorov-Smirnov test *** p < 0.001). *Center*: Time course for the average change in firing frequency upon nicotine injection for activated (red, n = 155) and inhibited (blue, n = 90) VTA DA neurons. *Right*: Average amplitude of the nicotine response for activated (red, mean = 33.75 ± 52 %) and inhibited DA neurons (blue, mean = −35.42 ± −2 %), compared to saline injection (grey, n = 147 and 86, respectively) (Wilcoxon test *** p < 0.001). **(C)** Analysis of the spontaneous activity of NB-labeled DA neurons that were either activated (red) or inhibited (blue) by the nicotine injection. *Left*: The two subpopulations of DA neurons did not display any significant difference in their firing rates (Wilcoxon test p = 0.054). However, neurons that were inhibited by nicotine displayed a basal percentage of spikes-within-burst (%SWB) higher than activated ones (Wilcoxon test * p = 0.04). *Right:* Interval between SWB (in ms) as a function of the length of the burst (from two to six action potentials). **D)** Localization of NB-labeled, nicotine-activated and -inhibited DA neurons, positioned on the Paxinos atlas at bregma - 3.3 mm. The subpopulation of nicotine-inhibited neurons had a more medial distribution in the VTA than the nicotine-activated subpopulation (Wilcoxon test *** p < 0.001), but neither antero-posterior (Wilcoxon test p = 0.4) nor dorso-ventral (Wilcoxon test p = 0.5) differences in their distribution were observed.

### The two subpopulations of DA neurons in the VTA, activated or inhibited by nicotine injection, have distinct axonal projections to the nucleus accumbens or the amygdala

The DA system is heterogeneous, and is increasingly thought about in terms of anatomically and functionally distinct subnetworks (Watabe-Uchida et al., 2012). DA neurons in the VTA have been reported to project to different terminal regions based on their localization along the mediolateral axis (Beier et al., 2019; 2015; Lammel et al., 2008), thus we next investigated whether these two subpopulations belong to anatomically distinct dopamine circuits by probing the nicotine-evoked responses of DA neurons with identified projection sites. To do so, green retrobeads (RB), a retrograde tracer, were injected either in the nucleus accumbens (NAc: shell and core) or in the amygdala nuclei (Amg: basolateral and central amygdala) (Figure S2.1). Two weeks later, spontaneous and nicotine-evoked activity of VTA DA neurons were recorded *in vivo* in anesthetized mice, and then all neurons were labeled with neurobiotin. Triple labeling immunofluorescence allowed us to confirm *post-hoc* the DA nature (TH+), projection site (RB+ / RB-), and position (NB+) of the recorded neurons (Figure 2A). We first noted that, in line with previous reports (Lammel et al., 2008), Amg-projecting DA neurons are located more medially in the VTA than NAc-projecting DA neurons (Figure S2.1). We identified 32 nicotine-activated and 17 nicotine-inhibited cells in mice with RB injected in the NAc, and 26 nicotine-activated and 26 nicotine-inhibited cells in mice with RB injected in the Amg (Figure 2B, E). For mice with RB injection in the NAc, 93% (28/30) of NAc-projecting VTA DA neurons (RB+, TH+) were activated by a nicotine injection, while 7% (2/30) of neurons were inhibited. In contrast, in these mice 79% (15/19) of DA neurons without evidence for projection to the NAc (RB-, TH+) were inhibited by a nicotine injection, and only 21% (4/19) of RB-neurons were activated (Figure 2C). For mice with RB injection in the Amg, 86% (19/22) of Amg-projecting VTA DA neurons (RB+, TH+) were inhibited by a nicotine injection, while 14% (3/22) were activated. Whereas, in these mice, DA neurons without evidence for projection to the Amg (RB-, TH+) were mainly activated by a nicotine injection (77%, 23/30), with 23% (7/30) of neurons inhibited (Figure 2E, F). Overall, these results indicate that the majority of VTA DA neurons activated by nicotine injection project to the NAc, whereas the majority of VTA DA neurons inhibited by nicotine project instead to the Amg. We next took advantage of this anatomical distinction to analyze their respective electrophysiological properties in *ex vivo* patch-clamp recordings on coronal VTA slices with NAc-projecting DA cells or Amg-projecting DA cells labeled with RB (Figure S2.2). Amg-projecting DA neurons demonstrated higher excitability (Figure S2.2B) than NAc-projecting DA neurons, but no difference in nicotine-evoked currents was found between these two populations (Figure S2.2D). These results indicate that these two VTA DA cell populations have different membrane properties, but do not markedly differ in the expression or function of nAChRs.

**Figure 2:**
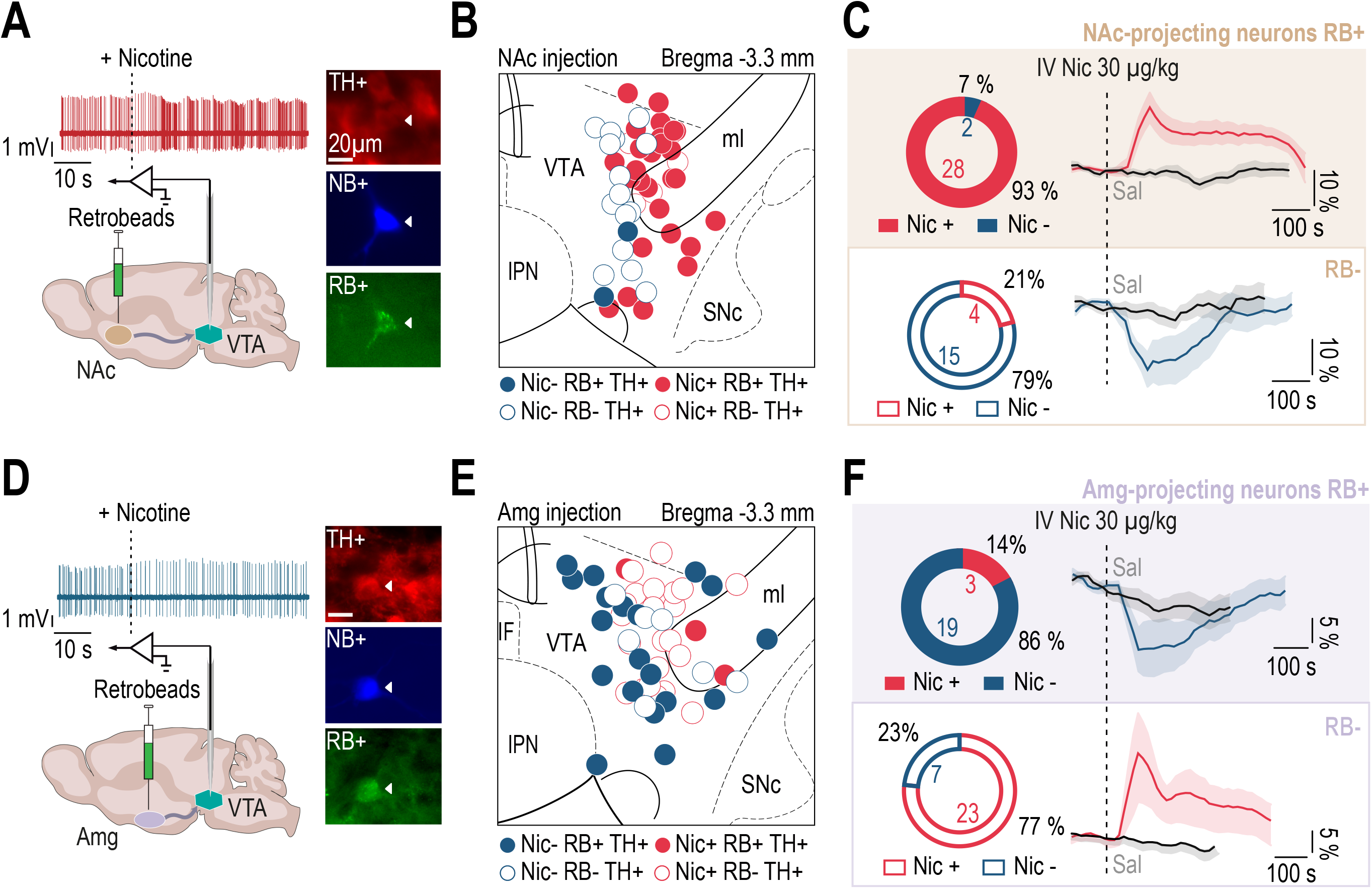
Nicotine evoked opposite responses in VTA DA neurons that project to the NAc or to the Amg. **(A)** Retrobeads (RB) were injected in the NAc, and three weeks later VTA DA neuron responses to an IV nicotine injection were recorded *in vivo,* and neurons were subsequently labeled with neurobiotin. DA neurons that project to the NAc were identified *post-hoc* by immunofluorescent co-labeling of TH, NB and RB. **(B)** Localization of NB-labeled DA neurons (NB+ TH+, n = 49) following RB injection into the NAc. RB+ and RB− neurons are represented by filled and empty circles, respectively. Red and blue colors denote nicotine-activated neurons (Nic+) and -inhibited neurons (Nic-), respectively. (RB+ Nic+, n = 28; RB+ Nic-, n = 2; RB− Nic+, n = 4; RB− Nic-, n = 15). **(C)** *Top:* Percentage and number of Nic+ (red) and Nic- (blue) cells among NAc-projecting DA neurons (RB+). Mean change in firing frequency of NAc-projecting DA neurons in response to an IV injection of nicotine (red) or saline (black). *Bottom*: Percentage and number of Nic+ (red) and Nic- (blue) cells in non RB-labeled neurons (RB-). Mean change in firing frequency of RB− DA neurons in response to an IV injection of nicotine (red) or saline (black). **(D)** In separate experiments, RBs were injected in the Amg, and two weeks later VTA DA neuron responses to an IV Nic injection were recorded *in vivo,* and neurons were subsequently labeled with neurobiotin. DA neurons that project to the Amg were identified *post-hoc* by immunofluorescent co-labeling of TH, NB and RB. **(E)** Localization of NB-labeled DA neurons (NB+ TH+, n = 52) following RB injection into the Amg. (RB+ Nic+, n = 3; RB+ Nic-, n = 19; RB− Nic+, n = 23; RB− Nic-, n = 7). **(F)** *Top:* Percentage and number of Nic- (blue) and Nic+ (red) among Amg-projecting DA neurons (RB+) or in RB− DA neurons of the VTA in mice injected in the Amg. Mean change in firing frequency of RB+ DA neurons in response to an IV injection of nicotine (red) or saline (black). *Bottom:* Percentage and number of Nic- (blue) and Nic+ (red) cells in non RB-labeled neurons (RB-). Mean change in firing frequency of RB− DA neurons in response to an IV injection of nicotine (red) or saline (black).

### The anxiogenic effect of nicotine injection involves β2 subunit-containing nAChRs in the VTA

We next asked whether these two distinct dopamine sub-circuits are associated with different behavioral outcomes after an acute injection of nicotine. Nicotine is known to have rewarding properties, through the activation of VTA DA neurons (Durand-de Cuttoli et al., 2018; Maskos et al., 2005; Tolu et al., 2013). Nicotine can also induce negative outcomes such as anxiety-like behaviors and stress-induced depressive-like states (Kutlu and Gould, 2015; Morel et al., 2017; Picciotto and Mineur, 2013). However, whether these negative effects are mediated by VTA DA neurons is unknown. We thus asked whether the anxiogenic effect of nicotine is mediated by VTA DA neurons, by analyzing mouse behavior in an elevated-O-maze (EOM). First, we found that mice injected with nicotine (intra-peritoneal, IP, 0.5 mg/kg) spent less time in the open arms of the EOM when compared to mice injected with saline, indicating that nicotine triggers anxiety-like behavior in mice (Figure 3A and Figure S3.1 for individual data). This anxiety-like phenotype was not related to a noticeable nicotine effect on locomotor activity (Figure S3.1). To provide evidence for a specific role of VTA neuron response to nicotine in this anxiogenic effect, we then locally infused nicotine into the VTA by a fixed implanted cannula, and found that, just as observed with the IP nicotine injection, these mice spent less time in the open arms of the EOM than those injected with saline (Figure 3B and Figure S3.1 for individual data). Local nicotine infusion in the VTA was therefore sufficient to trigger such anxiogenic effect. Finally, as nicotine-evoked responses have been shown to be mainly mediated by β2-containing nAChRs (β2*nAChRs) expressed on the soma of both DA and GABA neurons of the VTA (Tolu et al., 2013), we next assessed the role of β2*nAChRs within the VTA in mediating the anxiogenic effects of nicotine by using knock-out mice deleted for the nAChR β2 subunit (β2^−/−^ mice). Electrophysiological recordings of VTA DA neurons in β2^−/−^ mice demonstrated neither excitatory nor inhibitory responses to nicotine injection. In contrast, lentiviral re-expression of the β2 subunit restricted to the VTA of β2^−/−^ mice (β2^−/−^Vec mice), restored both excitatory and inhibitory responses to nicotine (Figure 3C). Finally, we found that β2^−/−^ mice injected or not with GFP expressing virus (see Figure S3F) were impervious to the anxiogenic effect of nicotine in the EOM, as nicotine did not reduce the time spent by β2^−/−^ mice in the open arms, but that this effect could be restored by re-expression of the β2 subunit within the VTA (β2^−/−^Vec, Figure 3D). Together, these results indicate that both the activation or inhibition of VTA DA neurons by nicotine, as well as the anxiogenic effects of nicotine in the EOM, require signaling through β2*nAChRs in the VTA. However, the present data do not allow us to conclude whether the anxiogenic effect of nicotine requires the activation of VTA DA neurons, their inhibition, or both types of response.

**Figure 3:**
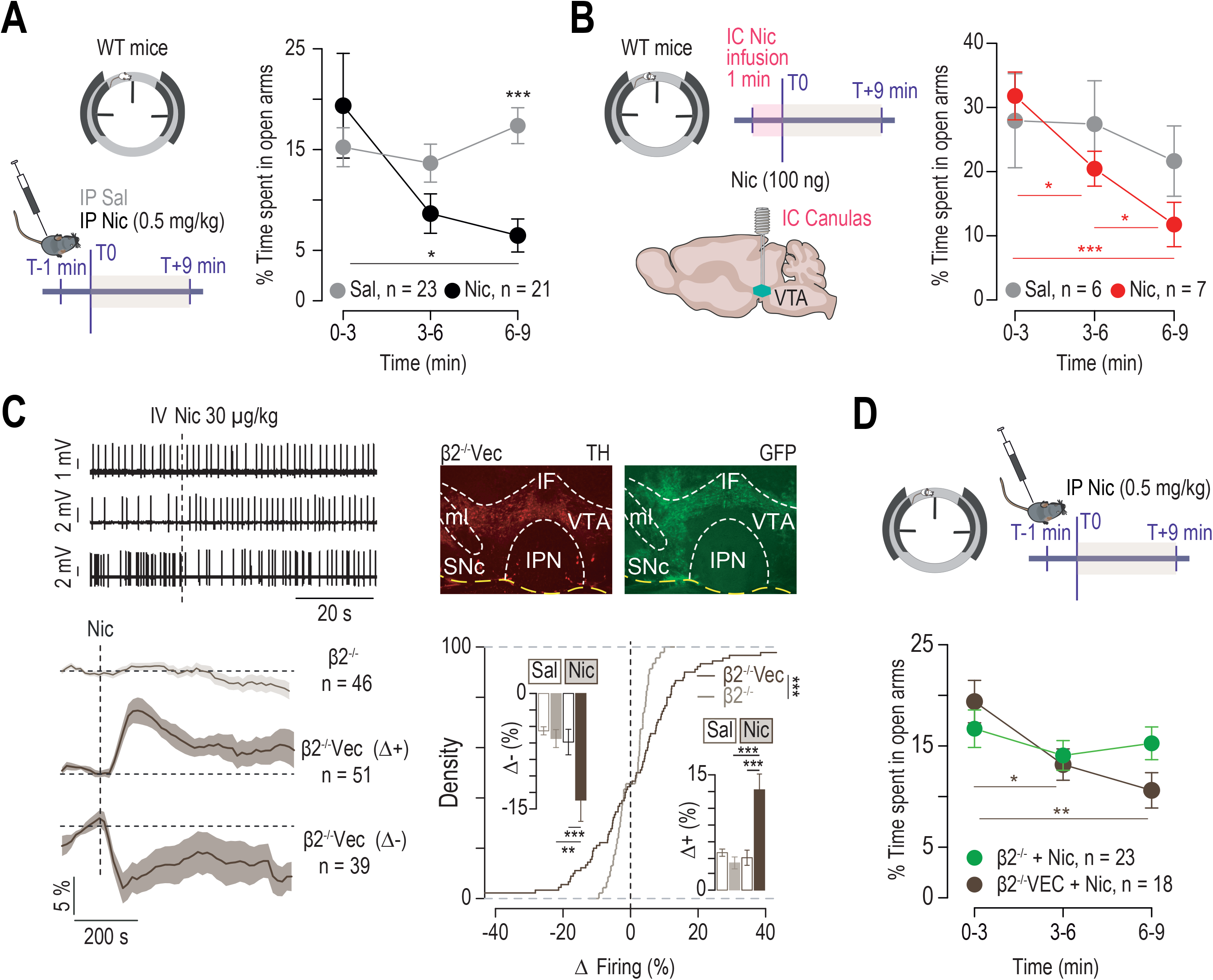
β2 subunit-containing nAChRs mediated both VTA DA neuron responses to nicotine injection and nicotine-induced anxiogenic effects. **(A)** Nicotine (Nic, 0.5 mg/kg) or saline (Sal) were injected intraperitoneally (IP) 1 minute before the 9-minute elevated O-maze (EOM) test. In wild-type (WT) mice (n = 21), nicotine injection decreased the time spent in the open arms of the EOM compared to the group injected with Sal (n = 23) (two-way RM ANOVA main effect of time F_(2,84)_ = 3.84, * p = 0.025, treatment x time interaction F_(2,84)_ = 5.37, ** p = 0.006; *post-hoc* Wilcoxon test with Bonferroni corrections * p (3 vs 9 minutes) = 0.03, p (3 vs 6 minutes) = 0.1, p (6 vs 9’) = 0.2; *post-hoc* Wilcoxon test Sal vs Nic at 9 minutes *** p < 0.001). **(B)** Mice implanted with intracranial guide cannulas were infused with either Sal or Nic (100 ng in 100 nl infusion) over 1 minute before the 9-minute EOM test. The Nic-injected group (n = 7) spent less time in the open arms over the test duration than the control group (n = 6) (two-way RM ANOVA main effect of time F_(2,22)_ = 12.48, *** p < 0.001, treatment x time interaction F_(2,22)_ = 9.66, *** p < 0.001; *post-hoc* Student’s t-test with Bonferroni corrections: *** p (3 vs 9 minutes) < 0.001, * p (3 vs 6 minutes) = 0.03, * p (6 vs 9 minutes) = 0.02; *post-hoc* Student’s t-test Sal vs Nic at 9 minutes, p = 0.054). **(C)** *Left*: Juxtacellular recording traces of VTA DA neurons in mice deleted for the β2 nAChR subunit (β2^−/−^) and in β2^−/−^ mice transduced with the β2 subunit-together with GFP-in the VTA (β2^−/−^Vec). Individual and mean responses (expressed as percentage of firing frequency variation) indicate that there were no nicotine-evoked responses in VTA DA neurons of β2^−/−^ mice (n = 46 cells from 12 mice), and that both nicotine-evoked activation (n = 51 cells from 18 mice, Δ+) and inhibition (n = 39 cells from 19 mice, Δ-) of VTA DA neurons were restored in β2^−/−^Vec mice. *Right*: Immunofluorescence for TH and GFP on β2^−/−^Vec mice. Cumulative distribution of nicotine-evoked response amplitude of VTA DA neurons in β2^−/−^ mice (n = 46 cells from 12 mice, grey) and β2^−/−^Vec mice (n = 90 cells from 24 mice, black) (Kolmogorov-Smirnov test ** p = 0.008). Bar plots show the maximum firing variation induced by nicotine (filled bars) and saline (unfilled bars) in the two groups. Nicotine injection did not alter the firing frequency of VTA DA neurons in β2^−/−^ mice, however it induced a significant increase (mean 12.45 ± 13.37) or decrease (mean −13.16 ± 16.31) in the firing frequency of VTA DA neurons in β2^−/−^Vec mice compared to saline (*** p < 0.001, *** p < 0.001) or β2^−/−^ mice (***p < 0.001, ** p = 0.005) (Wilcoxon paired test with Bonferroni corrections) **(D)** IP nicotine injection (0.5 mg/kg) in the EOM, for a control group (n = 23) of β2^−/−^ mice in which some were injected in the VTA with GFP (n = 6/23) and for β2^−/−^Vec mice (n = 18). Re-expression of β2 subunit in the VTA restored the nicotine-evoked anxiogenic effects in the EOM, which were absent in the β2^−/−^GFP mice (two-way RM ANOVA main effect of time F_(2,78)_ = 6.87, ** p = 0.002, treatment x time interaction F_(2,78)_ = 3.43, * p = 0.04; *post-hoc* Student’s t-test with Bonferroni corrections: ** p (3 vs 9 minutes) = 0.003, * p (3 vs 6 minutes) = 0.04, p (6 vs 9 minutes) = 0.2; *post-hoc* Student’s t-test β2^−/−^ and β2^−/−^GFP vs β2^−/−^Vec mice at 9 minutes, p = 0.06).

### The two projection pathways of DA neurons of the VTA underlie opposite functions in reinforcement and anxiety

To dissociate whether nicotine-evoked activation or inhibition of VTA DA neurons is necessary for the behavioral effects of nicotine, one would ideally need to isolate these responses in DA neurons, as well as in VTA GABA neurons, which also express nAChRs (Grieder et al., 2019; Tolu et al., 2013). Because activation and inhibition by nicotine are concomitant and not easily isolable from one another, and because the responses of VTA DA and GABA neurons to nicotine are also tightly linked (Tolu et al., 2013), we decided to mimic each response separately with an optogenetic approach. We used a calcium translocating channelrhodopsin (CatCh (Kleinlogel et al., 2011) for activation and a red-shifted cruxhalorhodopsin (Jaws (Chuong et al., 2014)) for inhibition (Figure S4.1). Dopamine transporter (DAT)- Cre mice, in which Cre recombinase expression is restricted to DA neurons without disrupting endogenous DAT expression (Turiault et al., 2007; Zhuang et al., 2005), were bilaterally injected in the VTA with CatCh (AAV5-flox-EF1a-hCatCh-YFP), Jaws (pXR5-CAG-flex-Jaws-eGFP), or a control enhanced yellow fluorescent protein (eYFP) with no opsin (AAV5-flox-EF1a-YFP, Figure S4.2A-B), and optical fibers were implanted either in the Basolateral Amg (BLA, Figure S4.2C) or in the NAc lateral shell (NAcLSh, Figure S4.2D) of these mice to restrict the effects of optogenetic stimulation to DA terminals within these regions.

We first examined the effects of these manipulations in the EOM test. Light-evoked inhibition of DA neuron terminals in the BLA reduced the percentage of time spent by Jaws-expressing mice in the open arms of the EOM, while no effect of the light stimulation was detected in control mice expressing only eYFP (Figure 4A and Figure S4.3A for individual data). Light-evoked activation of DA terminals in the BLA showed the opposite effect, increasing the percentage of time spent by CatCh-expressing mice in the open arms of the EOM compared to mice expressing eYFP (Figure 4B and Figure S4.3B for individual data). In contrast, light-evoked activation of DA neuron terminals in the NAcLSh of CatCh-expressing mice had no effect on the time spent in the open arms of the EOM compared to control mice expressing eYFP (Figure 4C and Figure S4.3C for individual data), and there was no noticeable effect of the light-stimulation on locomotor activity in any of these groups of DAT-Cre mice (Figure S4.3D-F). To determine whether the anxiogenic effect observed during inhibition of DA neuron terminals in the BLA was specific to the BLA nuclei, we compared groups of WT mice injected with either Jaws or GFP in the VTA and implanted with bilateral optical-fibers either in the BLA or in the central amygdala (CeA), which both receive DA input (Figure S4.3). We found that inhibiting VTA neuron terminals decreased the percentage of time spent in the open arms of the EOM when optical fibers were implanted in the BLA, but not when they were implanted in the CeA (Figure S4.4). Finally, to directly link nicotine-evoked inhibition of DA neurons to its anxiogenic effect, we reasoned that an optogenetic activation of the BLA-projecting terminals of VTA DA neurons would counteract the nicotine-evoked inhibition of this pathway, and should therefore abolish the anxiogenic effect of the drug. Thus, DAT-Cre mice expressing CatCh in the VTA were injected with IP nicotine one minute before the EOM test, and received light stimulation in the BLA throughout the 9-minute test. Light-evoked activation of BLA DA terminals during the EOM abolished the anxiogenic effect of nicotine injection in CatCh expressing mice when compared to eYFP expressing controls, which decreased the percentage of their time spent in the open arms after IP nicotine injection (Figure 4D).

**Figure 4:**
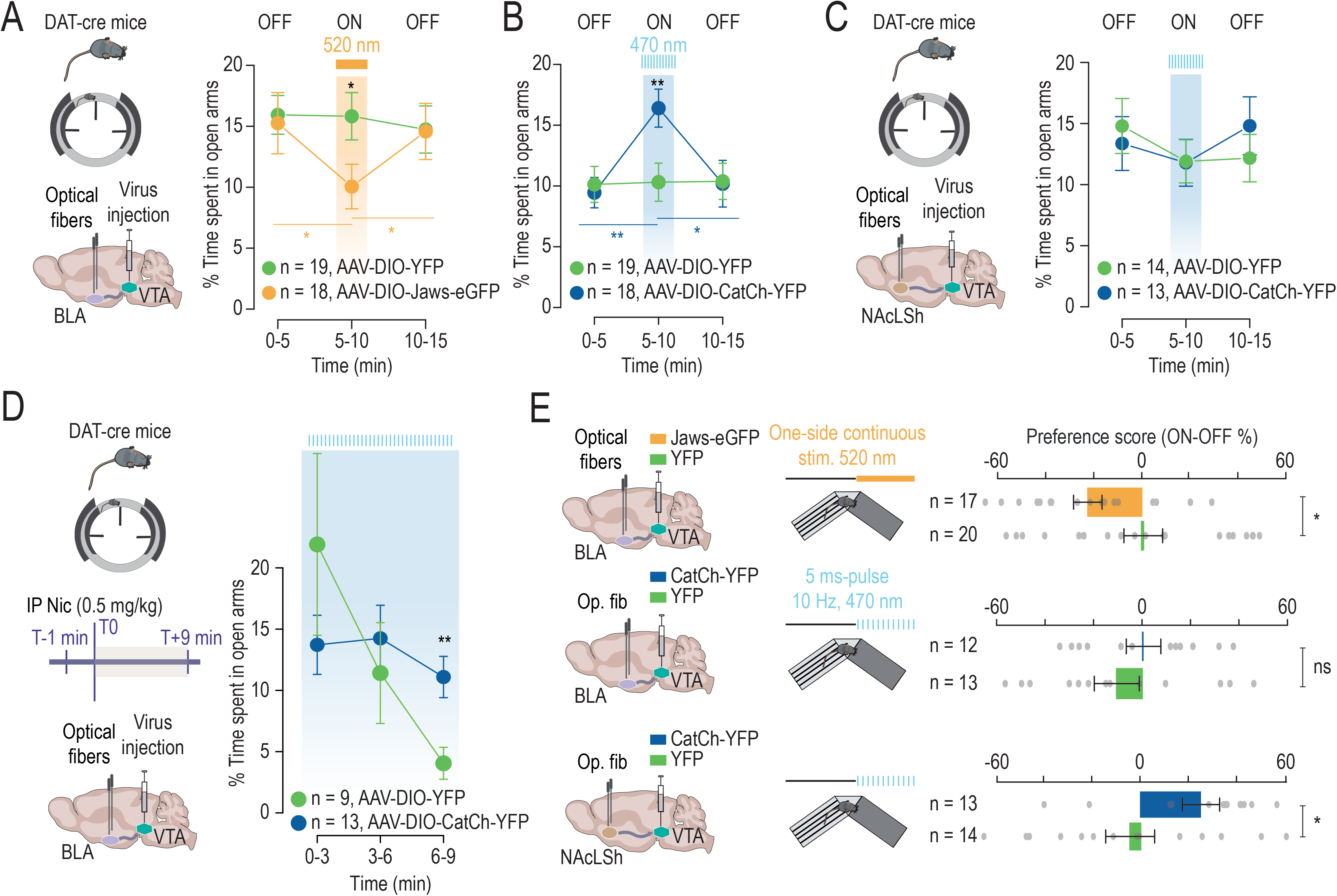
Selective optogenetic manipulation of VTA-BLA and VTA-NAcLSh pathways highlights the opposite effects of nicotine injection on VTA DA neurons. **(A)** Percentage of time spent in the EOM open arms for Jaws (orange, n = 18) and YFP control (green, n = 19) groups stimulated continuously at 520 nm over a 5-minute period (ON) in the BLA (two-way RM ANOVA time x opsin interaction F_(2,70)_ = 3.32, * p = 0.04; *post-hoc* Student’s t-test Jaws vs YFP: * p(ON) = 0.038; *post-hoc* Student’s t-test with Bonferroni corrections Jaws * p (5 vs 10 minutes) = 0.01, * p (10 vs 15 minutes) = 0.02) (**B)** Percentage of time spent in the EOM open arms for CatCh (blue, n = 18) and YFP control (green, n = 19) groups stimulated at 470 nm over a 5-minute period (ON) at 10 Hz, 5 ms-pulse in the BLA (Two-way ANOVA main effect of time F_(2,70)_ = 4.41, * p = 0.016, time x opsin interaction F_(2,70)_ = 4.43, * p = 0.015; *post-hoc* Student’s t-test YFP vs CatCh ** p(ON) = 0.009, *post-hoc* Student’s t-test with Bonferroni corrections CatCh ** p (5 vs 10 minutes) = 0.001; * p (10 vs 15 minutes) = 0.01) (**C**) Percentage of time spent in the EOM open arms for CatCh (blue, n = 13) and control YFP (green, n = 14) groups stimulated at 470 nm over a 5-minute period (ON) at 10 Hz, 5 ms-pulse in the NAcLSh, showing no difference between the CatCh and YFP groups. **(D)** Percentage of time spent in open arms in the EOM for CatCh (blue, n = 13) and YFP (green, n = 9) groups stimulated in the BLA throughout the test at 10 Hz, 5-ms light-pulse, after all receiving a nicotine IP injection (CatCh vs YFP: two-way RM ANOVA main effect of time F_(2,40)_ = 4.92, * p = 0.01, time x opsin interaction F_(2,40)_ = 3.74, * p = 0.03; one-way RM ANOVA YFP: F_(2,16)_ = 5.77, * p = 0.01; CatCh: F_(2,24)_ = 1.59, p = 0.6; *post-hoc* Student’s t-test YFP vs CatCh at 9 minutes ** p = 0.006). **(E)** Preference score in online place preference (OPP) paradigm for Jaws vs YFP groups bilaterally implanted in the BLA (*top*), for CatCh vs YFP groups bilaterally implanted in the BLA (*middle*), and for CatCh vs YFP groups bilaterally implanted in the NAcLSh (*bottom*). Animals were stimulated (continuously in the BLA, or at 10 Hz, 5-ms light-pulse, in the NAcLSh) over the 20 minute-test, and the preference score is defined by the % of time spent in the compartment where the animals are photo-stimulated compared to the compartment where they are not (ON-OFF). (*Top-down*) Optical inhibition of the VTA-BLA pathway (orange, n = 17) induced online place avoidance compared to the control group (YFP in green, n = 20) (Student’s t-test * p(ON-OFF) = 0.02). Mice with optical activation of the VTA-BLA pathway (blue, n = 12) did not display any difference in the OPP compared to the control group (YFP in green, n = 13) (Student’s t-test p(ON-OFF) = 0.5). Optical activation of the VTA-NAcLSh pathway (blue, n = 13) induced online place preference (Student’s t-test * p(ON-OFF) = 0.04) compared to the control group (YFP in green, n = 14).

We next investigated the effects of activating and inhibiting these two pathways on reinforcement using an online place preference paradigm (OPP). Activation of DA neuron terminals in the NAcLSh increased the preference score (percentage of time in ON – percentage of time in OFF) in mice expressing CatCh, indicating a rewarding effect of stimulating the VTA-NAcLSh pathway (Figure 4E). Light-evoked inhibition of DA neuron terminals in the BLA, however, reduced the preference score for the compartment where animals were photo-stimulated, and activation of these terminals showed no effect (Figure 4E). Together, these results argue for a functional dissociation of these two pathways: inhibition of Amg-projecting DA neurons evokes anxiogenic effects, while activation of NAc-projecting DA neurons mediates reward but has no effect on anxiety-like behavior. Furthermore, we show that the VTA-BLA DA pathway is the substrate for the anxiogenic effect of nicotine.

## Discussion

The VTA had long been perceived as a structure that broadly disseminates DA in the brain, with the different time courses of DA activity providing a phenomenological account for the functional involvement of DA neurons in different behavioral and neural processes (Schultz 2007). This temporal account of DA neuron function was gradually replaced or extended by the notion that the DA system, in particular the VTA, is divided into subpopulations of DA neurons, each associated with distinct appetitive, aversive, or attentional behaviors (Lammel et al., 2012). However, we are only beginning to appreciate how the functional activation/inactivation dynamics within these subpopulations impact behavioral and neural processes. Here, we show that activation and inhibition of VTA DA neurons appear concurrently as a consequence of nicotine injection. Furthermore, our results demonstrate that these activation/inhibition processes correspond to two anatomically and functionally distinct circuits, which mediate contrasting behavioral effects.

VTA DA neurons are known to be heterogeneous in their axonal projections, electrophysiological properties, and in several molecular features. They show for example striking differences in their expression of hyperpolarization-activated cyclic nucleotide-gated cation channels (HCN), the dopamine transporter (DAT), the dopamine receptor D2R, or vesicular glutamate transporters (VGLUTs) (Lammel et al., 2008; Margolis et al., 2008; Morales and Margolis, 2017). However, the role and the functional consequences of VTA DA neuron heterogeneity in behavior remain poorly understood. Here, we have demonstrated that nicotine evokes opposite responses in two distinct subpopulations of VTA DA neurons, those that project their axons to the NAc are activated by nicotine, while those that project to the Amg are inhibited. In addition to their functional and anatomical segregation, we found that these subpopulations display different bursting activities *in vivo*, and different excitabilities *in vitro*. However, they cannot be distinguished solely on the basis of their spontaneous firing pattern in anesthetized mice. Are there specific differences between these two neuronal populations, beside their projection sites, that would underlie their opposing responses to nicotine injection? NAc-projecting DA neurons exhibit smaller *I_h_* currents than BLA-projecting DA neurons, but have similar input resistances and capacitances (Ford et al., 2006), and similar expressions of DAT, D2R and TH (Su et al., 2019). We have previously reported that activated and inhibited DA cells react similarly to D2R agonist or antagonist injection *in vivo*, which is also consistent with similar D2R expression levels in the two neuronal populations (Eddine et al., 2015). Finally, there is no clear variation in nicotine-evoked currents in Amg or NAc-projecting DA cells, suggesting that nAChR expression does not differ markedly between these populations. Overall, we did not find striking evidence of intrinsic differences between NAc-projecting or Amg-projecting VTA DA neurons that could explain their differences in the nicotine-evoked responses. While differences may exist, it is also possible that the emergence of either nicotine-evoked activation or inhibition of these neurons arises from network dynamic. Within the VTA, nicotine directly activates both DA and GABA neurons, as they each express nAChRs (Klink et al., 2001; Tolu et al., 2013). In particular, β2*nAChRs of the VTA are key mediators of the positive reinforcing effects of nicotine, as previously shown by reexpressing β2 locally in the VTA of β2^−/−^ mice (Maskos et al., 2005; Tolu et al., 2013), or by rendering β2*nAChRs insensitive to nicotine using light (Durand-de Cuttoli et al., 2018). Here, we show that β2*nAChRs of VTA neurons are also required to evoke the inhibition of the VTA-Amg DA neuron subpopulation after systemic nicotine injection. Therefore, nicotine acting through β2*nAChRs activates VTA GABAergic interneurons and DA neurons that project to the NAc, while simultaneously inhibiting DA neurons projecting to the Amg. Several independent mechanisms may explain the inhibitory effect of nicotine on this subpopulation. These include (1) the activation of GABAergic interneurons that would, in turn, inhibit specifically this VTA DA neuron subpopulation; (2) inhibition through local DA release (Eddine et al., 2015), even though no difference in D2R-mediated inhibitory postsynaptic currents or in DA reuptake between NAc-projecting and BLA-projecting DA neurons has been reported (Ford et al., 2006); or (3) feedback inhibition. Indeed, subpopulations of DA neurons are embedded within distinct inhibitory networks resulting in specific feedback loops between VTA and NAc subregions (de Jong et al., 2019). NAc-projecting DA neurons could therefore inhibit DA neurons projecting to the Amg through an as yet unknown striatal relay.

Nicotine is highly reinforcing, but also generates aversive and anxiogenic effects at various doses (Balerio et al., 2006; Kutlu and Gould, 2015; Picciotto and Mineur, 2013; Wolfman et al., 2018). Thus, depending on the context, the exact same dose of nicotine can trigger anxiety and/or reinforcement. Aversion for high doses of nicotine and anxiety associated with nicotine withdrawal have been attributed to nicotinic and glutamatergic signaling in the habenulo-interpedoncular axis (Fowler et al., 2011; Frahm et al., 2011; Molas et al., 2017; Zhao-Shea et al., 2013). There is also evidence that nAChRs of neurons located in the Amg modulate depressive-like states (Mineur et al., 2016). However, a role for DA in aversion has also been proposed. D1R and D2R antagonists prevent conditioned-place aversion induced by an acute high-dose nicotine injection (Grieder et al., 2012), and β2*nAChRs were shown to be necessary for both the aversive and rewarding effects of nicotine using a subunit re-expression strategy in VTA DA and GABAergic neurons of β2^−/−^ mice (Grieder et al., 2019). However, the mechanism underlying these opposite effects of the drug has not yet been established. Here, we show that activation of β2*nAChRs of VTA neurons is necessary for nicotine to inhibit Amg-projecting DA neurons and induce anxiety-like behavior. This indicates that DA signaling is critically involved in the acute anxiogenic effect of nicotine and could also mediate aversion to nicotine. Moreover, injections of nicotine at the doses used in this study are known to be rewarding in different paradigms in mice, which has been attributed to VTA DA neuron activation (Durand-de Cuttoli et al., 2018; Maskos et al., 2005; Tolu et al., 2013). Overall, our study shows that the same intake of nicotine can induce a rewarding effect by activating the VTA-NAc dopamine pathway, and simultaneously signal a negative emotional state by inhibiting the VTA-Amg dopamine pathway.

Our findings emphasize the complex role of the DA system in not only positive but also negative motivational processes, and promote a more nuanced view of the effects of reinforcing doses of nicotine on VTA DA neurons. Opposing responses of DA neurons to drug exposure have also been observed with cocaine (Mejias-Aponte et al., 2015), ethanol (Doyon et al., 2013), and morphine (Margolis et al., 2014). Notably, the inhibition induced by opioids differs in BLA-projecting and NAc-projecting VTA DA neurons (Ford et al., 2006), suggesting that the behavioral effects of opioid drugs could also result from a specific pattern of inhibition in these two pathways. Since our results demonstrate that both rewarding and anxiogenic messages occur simultaneously upon nicotine exposure, and are conveyed by distinct subpopulations of VTA DA neurons, the question then arises as to how the concurrent engagement of two circuits with opposing messages could compete to produce nicotine reinforcement, and whether an imbalance between the two would lead to addiction. Indeed, this question may prove critical when it comes to medical strategies aimed at smoking cessation. While the optogenetic strategies used in this study are well suited to mimic the individual effects of a drug that also produces strong and synchronized neuronal activity, the translational value of these effects is perhaps not to be sought in the specific activation or inhibition of a given neuronal pathway, but rather in the functional imbalance that this creates between the target structures of VTA neurons. Nevertheless, a detailed understanding of the multiple pathways engaged in nicotine-evoked responses and of their respective behavioral contributions can still help us understand the mechanisms leading to nicotine addiction. In this respect, the activation and inhibition processes which appear in VTA DA neurons as a consequence of systemic nicotine injection call for further mechanistic studies, but we show that they correspond to discrete neuronal circuits and that they mediate distinct behavioral effects, both of which are relevant to the understanding of addiction.

## Acknowledgements

We thank Lauren Reynolds for critical reading of the manuscript. We are grateful to France Lam and the imaging platform facility (IBPS), the animal facilities (IBPS), Victor Gorgievski for behavioral data acquisition. We are grateful to Mélissa Desrosiers, Camille Robert and Paris Vision Institute AAV production facility for viral production and purification.

This work was supported by the Centre National de la Recherche Scientifique CNRS UMR 8246, INSERM U1130, the Foundation for Medical Research (FRM, Equipe FRM DEQ2013326488 to P.F), FRM FDT201904008060 (to SM), the French National Cancer Institute Grant TABAC-16-022 et TABAC-19-020 (to P.F.), French state funds managed by the ANR (ANR-16 Nicostress to PF, ANR-19 Vampire to FM), The LabEx Bio-Psy (to P.F and Doctoral Fellowship to CNG). PF and UM are members of LabEx Bio-Psy.

## Author contributions

C.N., F.M. and P.F. designed the study. C.N., F.M. and P.F. analyzed the data. C.N. and F.M. performed *in vivo* electrophysiological recordings. S.T. and S.V. contributed to *in vivo* electrophysiological recordings. S.M. performed *ex vivo* patch-clamp recordings and data analyses. T.L.B. contributed to *in vivo* electrophysiological data analyses. C.N. performed injections, fiber and cannula implantations. C.N. and I.C performed behavioral experiments. S.M. and B.H. contributed to behavioral experiments. R.D.C and A.M contributed to optogenetic experiments. C.N and S.M. performed immunostaining experiments. D.D., S.P. and U.M. provided viruses. U.M. provided ACNB2 KO mice. C.N., J.P.H., A.M., F.M. and P.F. wrote the manuscript.

## Data and materials availability

All the data are available from the corresponding authors upon request, including the R code routines.

## Supplementary Figures

**Figure S1.**
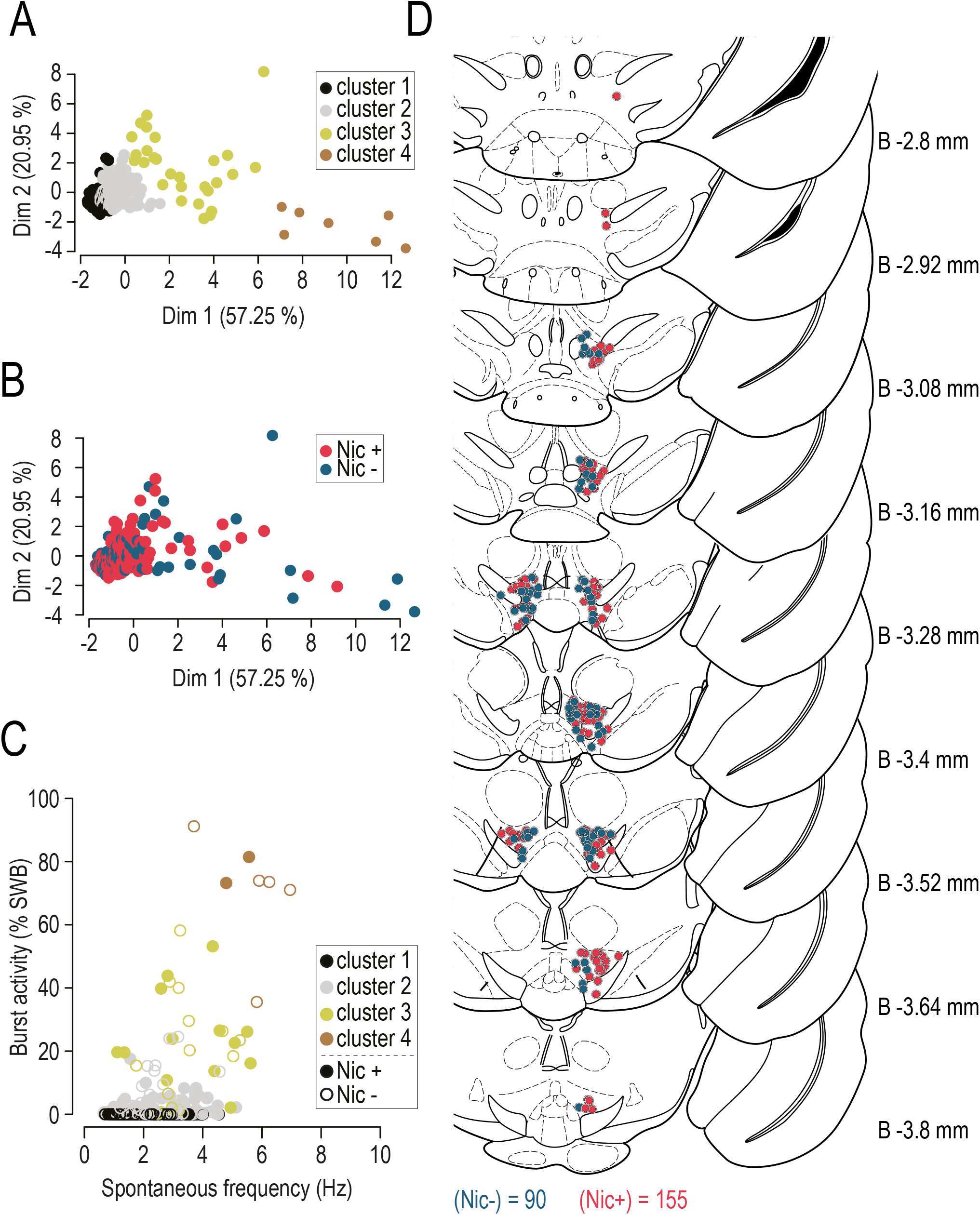
**(A-B**) Principal component analysis (PCA) of the spontaneous activity of VTA DA neurons labeled in vivo after juxtacellular recordings. First (axis Dim 1) and second (axis Dim 2) components are represented. Based on their basal activity, neurons are distributed in 4 clusters (cluster 1, n = 77; cluster 2, n = 132; cluster 3, n = 28; cluster 4, n = 7) (**A**), with nicotine-activated (in red, n = 154) and nicotine-inhibited (in blue, n = 90) neurons found in each of these clusters (**B**). **(C)** Mean firing frequency (Hz) as a function of percentage of spikes within a burst (%SWB) for the 4 clusters found with the PCA analysis, with the existence of nicotine-activated (cluster 1, n = 52; cluster 2, n = 85; cluster 3, n = 15; cluster 4, n = 2) and nicotine-inhibited (cluster 1, n = 25; cluster 2, n = 47; cluster 3, n = 13; cluster 4, n = 5) DA neurons in all of them. **(D)** Localization of VTA DA neurons labeled *in vivo* after juxtacellular recordings. Neurons are color-coded according to their responses to nicotine injection (activated in red, n = 155 and inhibited in blue, n = 90), and positioned according to the antero-posterior axis on the Paxinos atlas from Bregma - 2.8 to −3.8 mm.

**Figure S2.1.**
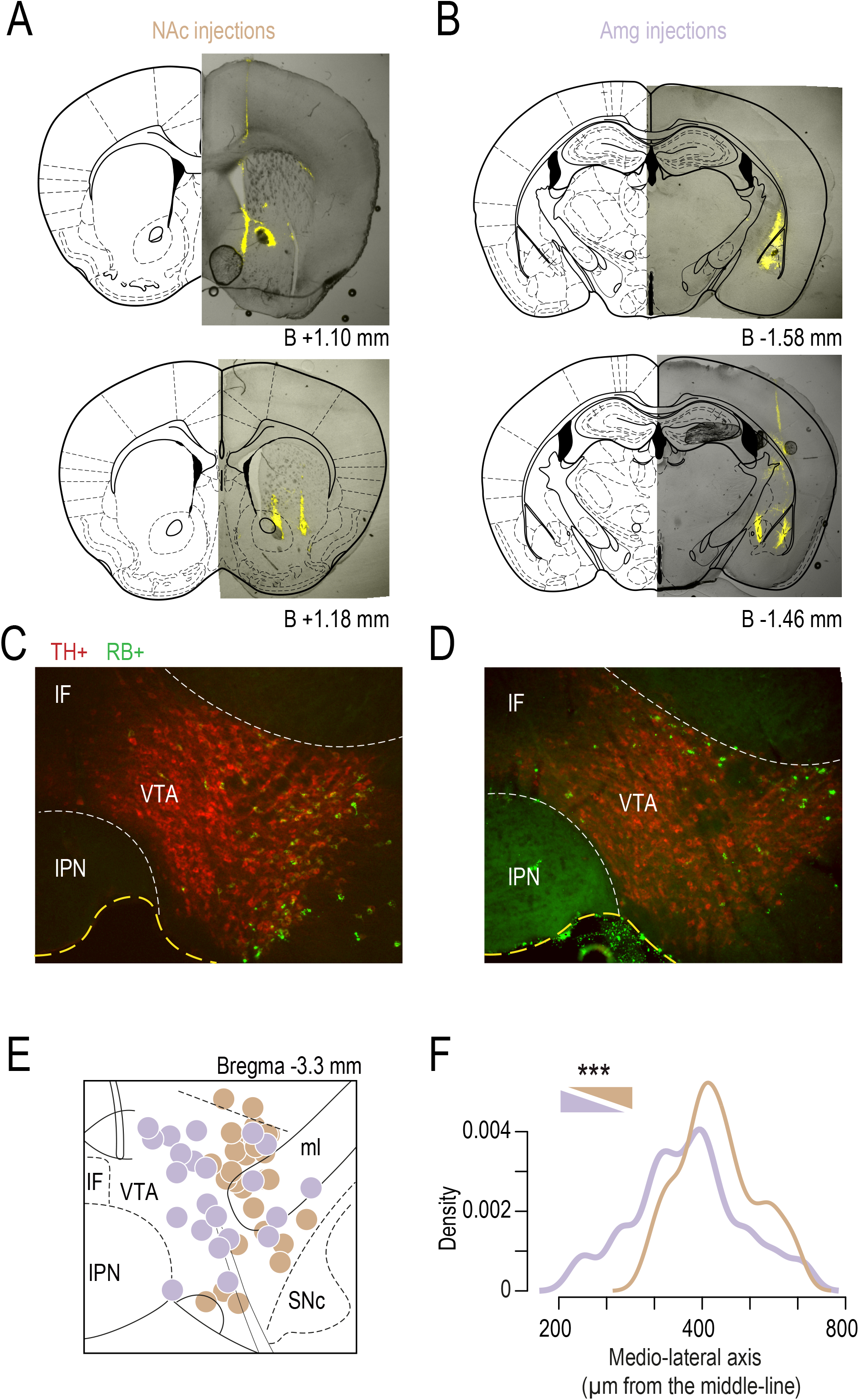
**(A, B)** Examples of NAc-injection sites **(A)** and Amg-injection sites **(B)** with retrobead (RB) tracer (in yellow) reported onto Paxinos atlas slices. **(C, D)** Examples of immunohistofluorescence analysis of VTA slices (TH+, red) revealing neuronal soma containing RB (RB+, green), after injection of the retrobeads into the NAc **(C)** or into the Amg **(D)**. **(E) A** Paxinos atlas slice at 3.3 mm from bregma onto which Neurobiotin-filled cell bodies of each recorded neuron were positioned. **(F)** Analysis of the medio-lateral distribution of recorded DA neurons of the VTA (shown as density) projecting either to the NAc (n = 30, gold) or to the Amg (n = 22, purple) revealed that Amg-projecting neurons are located more medially in the VTA than NAc-projecting neurons (Wilcoxon test, *** p < 0.001). IF: *interfascicular nucleus*; IPN: *interpeduncular nucleus*; ml: *medial lemniscus*; SNc: *substantia nigra pars compacta*

**Figure S2.2.**
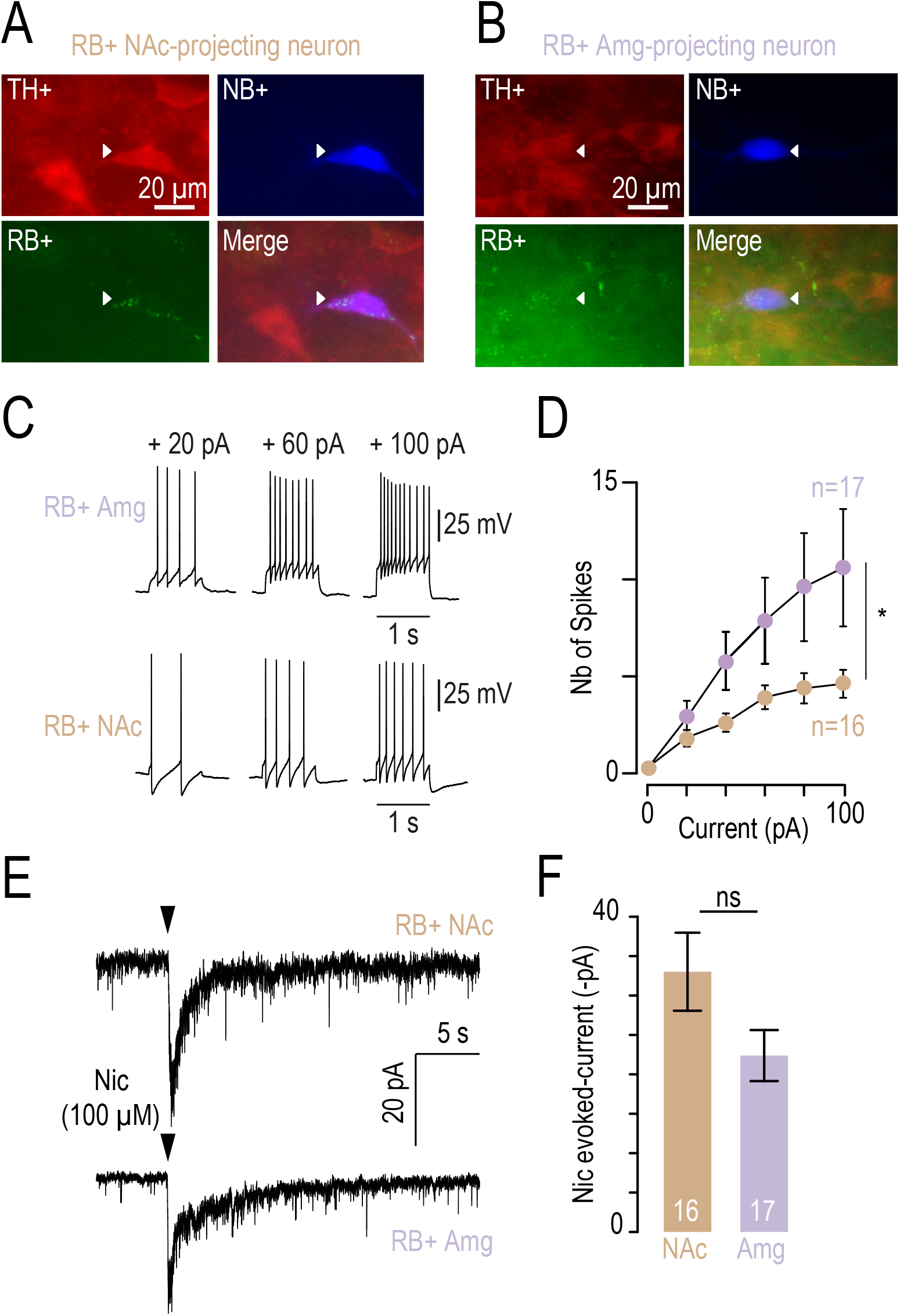
**(A)** Immunohistofluorescence analysis of a NAc-projecting neurons of the VTA labeled (neurobiotin, NB+) after patch-clamp recording: confirmed as DA (TH+) and NAc-projecting (retrobeads, RB+). **(B)** Immunohistofluorescence analysis of an Amg-projecting neuron of the VTA labeled (NB+) after patch-clamp recording: confirmed as DA (TH+) and Amg-projecting (RB+). **(C)** Firing of NAc-projecting and Amg-projecting DA neurons of the VTA after injection of currents (20, 60 and 100 pA). **(D)** Higher excitability of Amg-projecting (n = 17, purple) compared to NAc-projecting DA neurons (n = 16, gold) (two-way RM ANOVA main effect phenotype F_(1,26)_ = 4.96, * p = 0.035, current F_(4,104)_ = 15.97, *** p < 0.001, current x phenotype interaction F_(4,104)_ = 13.78, *** p < 0.001). **(E)** Nicotine-evoked currents (local puff 100 μM) in RB+-identified, NAc- or Amg-projecting VTA DA neurons recorded in brain slices (whole-cell voltage-clamp mode −60 mV). **(F)** Mean currents evoked by nicotine in either NAc-projecting (n = 16, gold, 33.0 ± 19.8 pA) or Amg-projecting (n = 17, purple, 22.4 ± 13.3 pA) VTA DA neurons are not statistically different (Student’s t-test p = 0.08).

**Figure S3.**
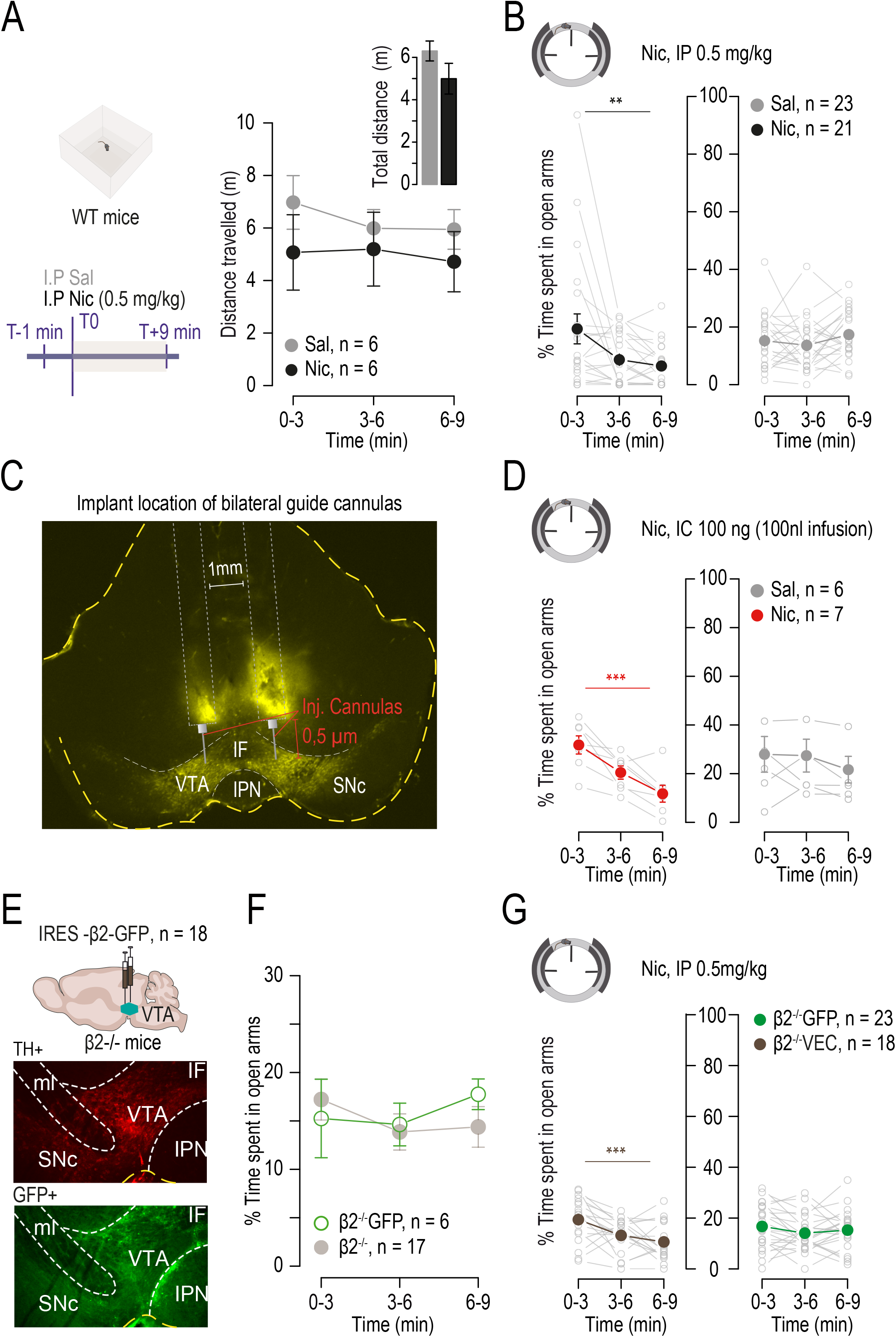
**(A)** Locomotor activity, measured in a square open field (OF), of mice receiving an intraperitoneal (IP) injection of either saline (IP Sal, grey, n = 11) or nicotine (IP Nic 0.5 mg/kg, black, n = 7). Mice did not display any statistically significant difference in the distance traveled over time (pooled each 3 minutes, two way RM ANOVA no time, no treatment or interaction effect p > 0.05) or in the total distance traveled during 9 minutes (presented as bar plot, Student’s t-test p > 0.05) **(B)** EOM detailed for WT mice tested with Sal (grey, n = 23) or Nic (black, n = 21) (one-way RM ANOVA Nic: F_(2,40)_ = 5.18, ** p = 0.01; Sal: F_(2,44)_ = 1.65, p = 0.2) after IP injection. Mean scores are represented in bold and color, and individual scores with empty grey dots. **(C)** Example of *post-hoc* verification of intracranial guide cannula implantations in WT mice. Bilateral injection cannulas (0.5 μm longer than the guide cannulas) are inserted on the day of the experiment, for local infusion into the VTA. TH labeling is shown in yellow. **(D)** Detailed scores of wild-type mice in the EOM after intracranial infusion of Sal (grey, n = 6) or Nic (red, n = 7) 1 mg/mL (one-way RM ANOVA Nic: F_(2,12)_ = 26.11, *** p <.001; Sal: F_(2,10)_ = 0.01, p = 0.99) 1 minute before the test. Mean scores are represented in bold and color, and individual scores in empty grey dots. **(E)** Schematic of β2 subunit re-expression by lentiviral vectorization in the VTA of β2^−/−^ mice. Lentivirus encoding either pGK-β2-IRES-GFP (β2^−/−^Vec) or pGK-GFP (β2^−/−^GFP) as a control were injected into the VTA. Example of immunohistofluorescence analysis of a β2^−/−^Vec mouse brain labeled for TH (red) and GFP (green). **(F)** The percent of time spent in the open arms of the EOM in β2^−/−^GFP animals (n = 6, green) and β2^−/−^ mice (n = 17, grey) are not different (two-way RM ANOVA no time, no treatment or interaction effect p > 0.05). **(G)** Detailed scores in the EOM test of β2^−/−^ mice (n = 23, green) and β2^−/−^Vec mice (n = 18, brown), after an injection of Nic 0.5 mg/kg (one-way RM ANOVA β2^−/−^Vec: F_(2,34)_ = 8.65, *** p < 0.001; β2^−/−^: F_(2,44)_ = 1.08, p = 0.3) 1 minute before the test. Mean scores are represented in bold and color, and individual scores in empty grey dots. IF: *interfascicular nucleus*; IPN : *interpeduncular nucleus*; SNc : *substantia nigra pars compacta;* ml : *medial lemniscus*; SNc : *substantia nigra pars compacta.*

**Figure S4.1.**
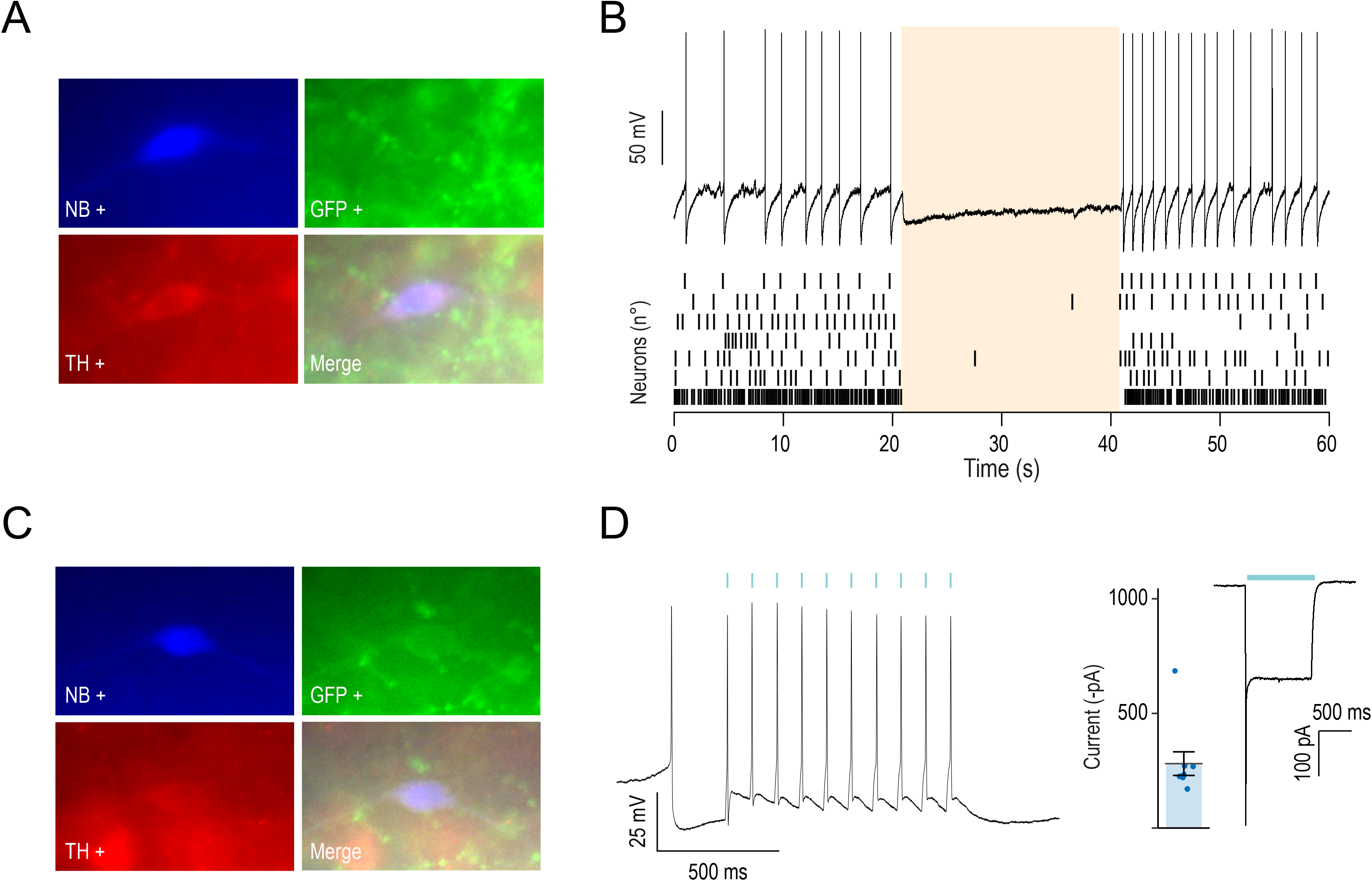
**(A)** Immunohistofluorescence analysis of VTA DA neurons after patch-clamp recordings in mice injected with AAV-Ef1α-DIO-Jaws-eGFP into the VTA. Neurobiotin (NB, blue), tyrosine hydroxylase (TH, red), YFP (green). **(B)** Example of a recording trace of a VTA DA neuron during continuous light stimulation (20 s, 520 nm) and raster plot of action potential showing light-induced inhibition in Jaws-expressing DA neurons (n = 7). **(C)** Immunohistochemistry of VTA DA neurons (NB, blue; TH, red; YFP, green) after patch-clamp recording in mice injected with AAV-DIO-CatCh-YFP into VTA. **(D)** Example of recording trace of a DA neuron of the VTA during light stimulation (10 Hz, 5-ms pulse, 470 nm) and light-evoked inward current in DA neurons expressing CatCh. Mean light-evoked current in seven DA neurons.

**Figure S4.2.**
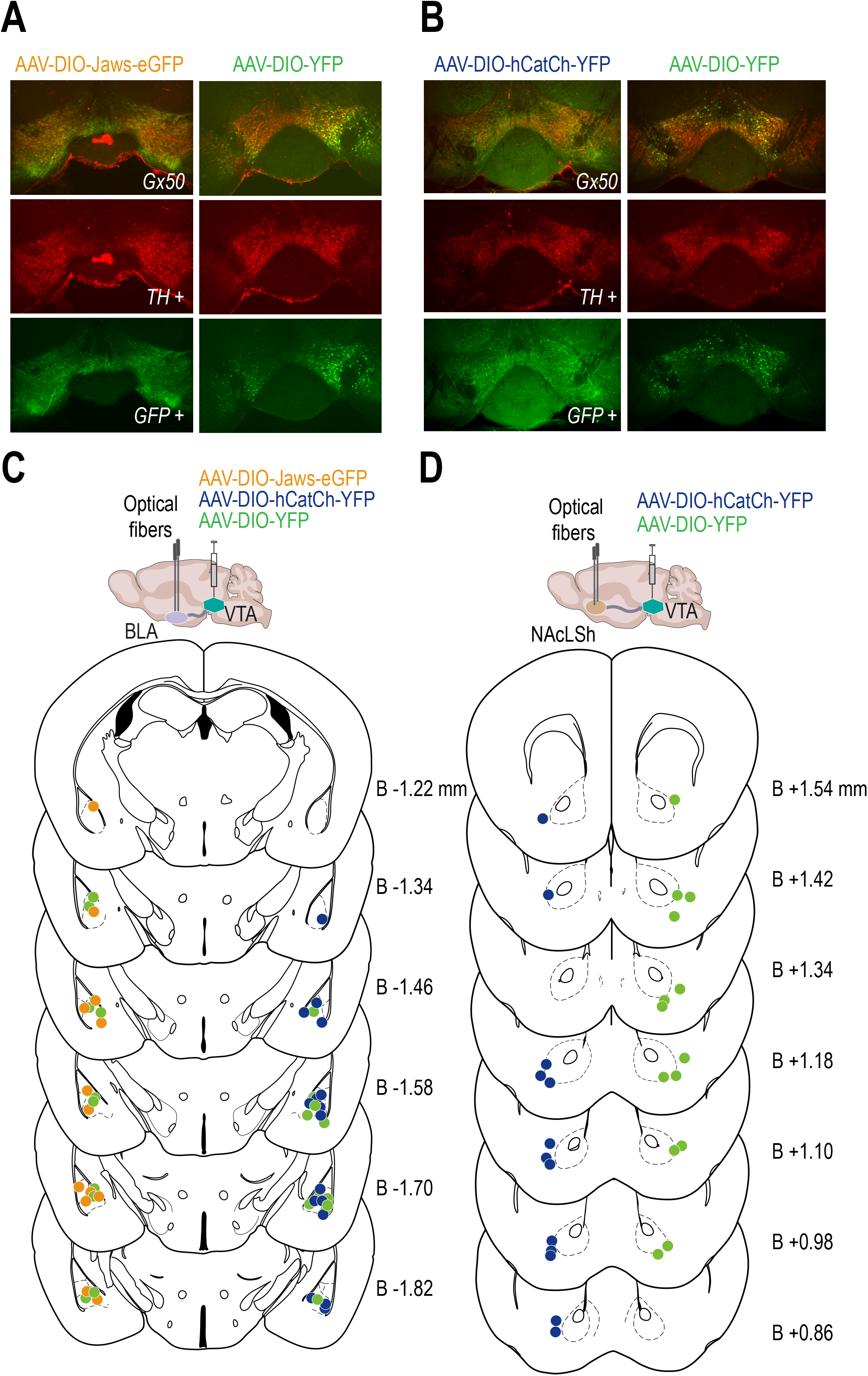
**(A)** Immunohistofluorescence analysis of VTA slices after AAV-DIO-Jaws-eGFP and AAV-DIO-YFP injections into VTA. **(B)** Immunohistofluorescence analysis of VTA slices after AAV-Ef1α-DIO-hCatCh-YFP and AAV-Ef1α-DIO-YFP injections into the VTA. **(C)** Example of *post-hoc* verification of the fiber implantation into the basolateral Amg (BLA) of mice used in optogenetic experiments, injected either with AAV-Ef1α-DIO-Jaws-eGFP (orange dots indicate the location of one hemisphere fiber tip that was verified, n = 13, left side of the slices) and its YFP control (green dots, n = 10, left side of the slices) or with AAV-Ef1α-DIO-hCatCh-YFP (blue dots, n = 16, right side of the slices) and its YFP control (green dots, n = 11, right side of the slices). The optical fibers were positioned onto Paxinos atlas slices from bregma −1.22 to −1.82 mm. **(D)** *Post-hoc* verification of the fiber implantation into the NAc lateral shell (NAcLSh) of mice used in optogenetic experiments, injected either with AAV-Ef1α-DIO-hCatCh-YFP (blue dots, n = 13, left of the slices) and its YFP control (green dots, n = 14, right of the slices). The optical fibers were positioned onto Paxinos atlas slices from bregma + 0.86 to + 1.70 mm.

**Figure S4.3.**
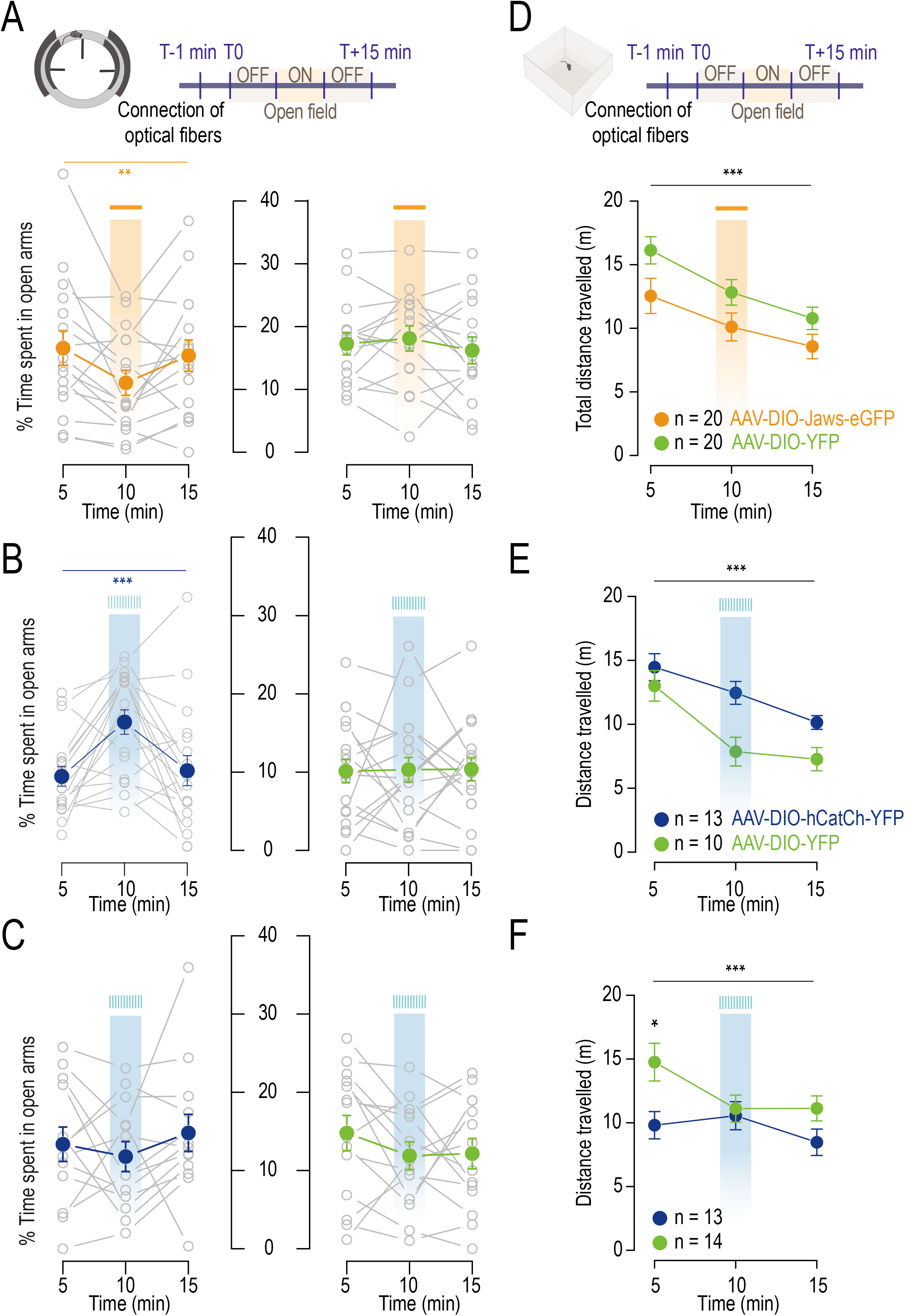
**(A)** Time spent in open arms of the EOM by mice injected with AAV-Ef1α-DIO-Jaws-eGFP (n = 18, orange, one-way RM ANOVA F_(2,34)_ = 5.28, ** p = 0.01) and AAV-Ef1α-DIO-YFP (n = 19, green, one-way RM ANOVA F_(2,36)_ = 0.32, p = 0.7) in the VTA, with (ON) or without (OFF) light-evoked inhibition of DA axon terminals in the BLA (the test lasts 15 minutes, with continuous photostimulation at 520 nm during a 5-minute ON period). Mean scores are represented in bold and color, and individual scores in grey. **(B)** Time spent in open arms of the EOM by mice injected with AAV-Ef1α-DIO-hCatCh-YFP (n = 18, blue, one-way RM ANOVA F_(2,34)_ = 9.27, *** p < 0.001) and AAV-Ef1α-DIO-YFP (n = 19, green, one-way ANOVA F_(2,36)_ = 0.01, p = 0.99) in the VTA, with (ON) or without (OFF) light-evoked activation of DA axon terminals in the BLA (10 Hz photostimulation at 470 nm, 5-ms pulse, during a 5-minute ON period). Mean scores and individual scores are shown as in (A). **(C)** Time spent in open arms of the EOM by mice injected with AAV-Ef1α-DIO-hCatCh-YFP (n = 13, blue, one-way RM ANOVA F_(2,24)_ = 0.61, p = 0.55) and AAV-Ef1α-DIO-YFP (n = 14, green, one-way RM ANOVA F_(2,26)_ = 1.47, p = 0.25) in the VTA, during (ON) or out (OFF) light-evoked activation of DA axon terminals in the NAcLSh (10 Hz photostimulation at 470 nm, 5-ms pulses, during a 5-minute ON period). Mean scores and individual scores are presented as previously described. **(D)** A square open field (OF) paradigm was conducted in the three paired groups of animals to assess their locomotor activities: Jaws- and YFP-injected mice implanted in the Amg (two-way RM ANOVA, main effect epoch F_(2,76)_ = 44.27, *** p < 0.001, no opsin or interaction effect). **(E)** CatCh- and YFP-injected mice implanted in the BLA (two-way RM ANOVA, main effect epoch F_(2,42)_ = 25.17, *** p < 0.001, no opsin or interaction effect). **(F)** CatCh- and GFP-injected mice implanted in the NAcLSh (two-way RM ANOVA, main effect epoch F_(2,50)_ = 14.27, ***p < 0.001, epoch x opsin interaction F_(2,50)_ = 4, * p = 0.02, *post-hoc* Wilcoxon test YFP vs CatCh at 5 minutes * p = 0.04). Animals showed no difference in terms of locomotor activity between the two groups or as a function of the stimulation period within a group.

**Figure S4.4.**
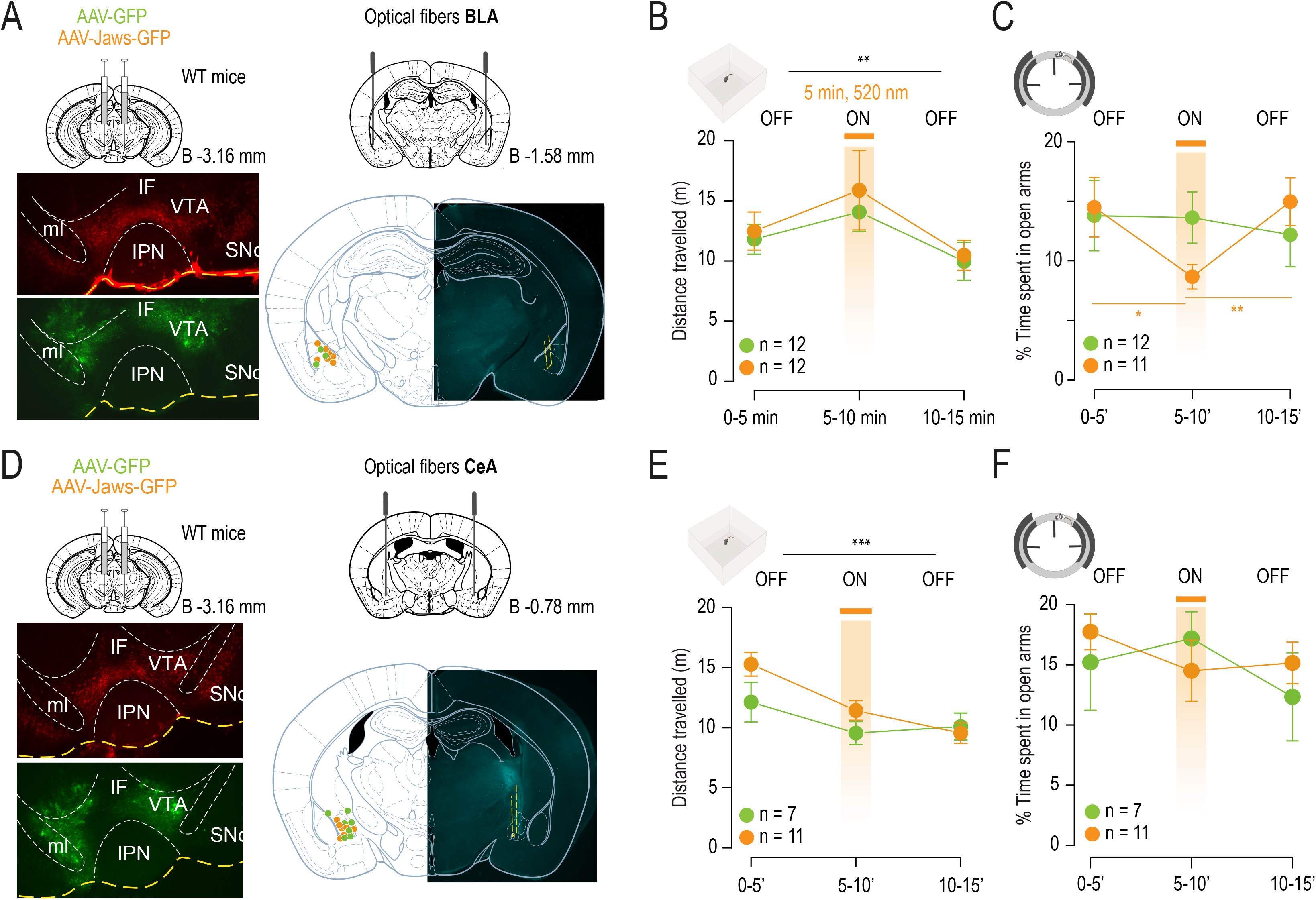
**(A)** (*Left*) Representative immunohistofluorescence analysis of VTA slices for Jaws expression (GFP: green labeling, TH: red labeling). (*Right*) Optical fibers were implanted in the basolateral Amg (BLA, bregma −1.61; lateral 3.18; ventral 4.7 mm) of wild-type (WT) mice injected either with AAV-CAG-Jaws-GFP (orange dots represent the location of one hemisphere fiber tip that was verified, n = 7) or with AAV-CAG-GFP control (green dots, n = 3) into the VTA. **(B)** Mice implanted in the BLA were tested for any difference in locomotor activity between groups in the open field (OF). The test lasted 15 minutes and consisted of a 5-minute period of photostimulation (continuous at 520 nm) in between two non-stimulation periods (OFF-ON-OFF). During both OFF- and ON-periods, the groups did not present any statistically significant difference in the distance traveled in the OF (two-way RM ANOVA main effect epoch F_(2,44)_ = 5.89, ** p = 0.005, no opsin or interaction effect*, post-hoc* Student’s t-test with Bonferroni corrections, p > 0.05). **(C)** Inhibiting BLA axon terminals by photostimulation of inhibitory opsin Jaws during the EOM task induced a decrease in the time spent by the mice in the open arms compared to the control group (two-way RM ANOVA, epoch x opsin interaction F_(2,42)_ = 3.44, * p = 0.04, *post-hoc* Student’s t-test p (ON Jaws vs GFP) = 0.056; *post-hoc* Student’s t-test with Bonferroni corrections Jaws *p (5 minutes vs 10 minutes) = 0.01; ** p (10 minutes vs 15 minutes) = 0.005). **(D)** (*Left*) Representative immunohistofluorescence analysis of VTA slices for Jaws expression (GFP: green labeling, TH: red labeling). (*Right*) Optical fibers were bilaterally implanted in the central amygdala (CeA) of another group of WT mice injected with either AAV-CAG-Jaws-GFP or with AAV-CAG-GFP control into the VTA. Orange dots and green dots represent the location of one hemisphere fiber tip that was verified in animals expressing Jaws (n = 7) and in control animals expressing GFP (n = 7), respectively. **(E)** Inhibiting CeA terminals by photostimulation did not produce any statistically significant difference in locomotor activity in the OF test between the Jaws- and GFP-expressing groups (two-way RM ANOVA main effect epoch F_(2,32)_ = 23.11, *** p < 0.001, epoch x opsin interaction F_(2,32)_ = 3.8, * p = 0.03, no opsin effect, *post-hoc* Student’s t-test with Bonferroni corrections, p > 0.05). **(F)** The two groups did not display any difference in time spend in the open arm of the EOM test either (two-way RM ANOVA, epoch x opsin interaction F_(2,32)_ = 3.67, * p = 0.04, *post-hoc* Student’s t-test with Bonferroni corrections p > 0.05). IF: *interfascicular nucleus*; IPN: *interpeduncular nucleus*; ml: *medial lemniscus*; SNc: *substantia nigra pars compacta*

## Materials and methods

### Animals

Wild-type C57Bl/6Rj (Janvier Labs, France), ACNB2 KO (β2^−/−^) and DAT^iCRE^ (DAT-Cre) male mice, weighing 25-35 grams, were used for all experiments. β2^−/−^ mice were generated using standard homologous recombination procedures. Founders were backcrossed onto a C57BL/6J background for a least 20 generations and bred on site. DAT^iCRE^ mice were provided by François Tronche’s team (IBPS Paris, France). They were bred on site and genotyped as described (Turiault et al., 2007).

Mice were kept in an animal facility where temperature (20 ± 1°C) and humidity were automatically monitored, and a circadian light-dark cycle of 12/12 hours was maintained. All experiments were performed on 8-to-16-week-old mice. All experiments were performed in accordance with the recommendations for animal experiments issued by the European Commission directives 219/1990, 220/1990 and 2010/63, and approved by Sorbonne University.

### AAV production

AAV vectors were produced as previously described (Khabou et al., 2018) using the co-transfection method, and purified by iodixanol gradient ultracentrifugation (Choi et al., 2007). AAV vector stocks were titrated by quantitative PCR (qPCR) (Aurnhammer et al., 2012) using SYBR Green (Thermo Fischer Scientific). Lentiviruses were prepared as previously described (Maskos et al., 2005; Tolu et al., 2013), with a titer of either 380 ng of p24 protein per μl or 764 ng/μl for the AChR β2-expressing vector, and 150 ng of p24 protein per μl or 361 mg per 2 μl for GFP-expressing vector.

### Drugs

The solution of nicotine (Nic) used for all experiments is a nicotine hydrogen tartrate salt (Sigma-Aldrich, USA). For juxtacellular recordings, we performed an intravenous injection of nicotine at a dose of 30 μg/kg (4.16 mg/kg, free base) or saline water (H_2_O with 0.9 % NaCl). For the behavioral test, in elevated O-maze (EOM) or open-field (OF), mice were injected intra-peritoneally with nicotine at 0.5 mg/kg 1-minute prior to the test. For intra-cranial (IC) experiments in EOM, saline solution or 100ng of Nic tartrate were infused in 100 nl/injection over 1 minute before the beginning of the test. All solutions were prepared in the laboratory.

### Retrobeads injection

Green fluorescent retrobeads (RB) tracers (LumaFluor Inc., Naples, FL) were injected (200 nl per site, 0.1 μl/min) in wild-type (WT) animals either in the NAc (NAc lateral shell NAcLatSh : bregma + 1.45 mm; lateral 1.75 mm; ventral 4.0 mm; NAc medial shell : bregma + 1.78 mm; lateral 0.45 mm; ventral 4.1 mm; NAc core : bregma 1.55 mm; lateral 1.0 mm; ventral 4.0 mm) or in the Amg (BLA : bregma - 1.61 mm; lateral 3.18 mm; ventral 4.7 mm; CeA : bregma - 0.78 mm; lateral 2.3 mm; ventral 4.8 mm) with a 10 μl Hamilton syringe (Hamilton) coupled with a polyethylene tubing to a 36G injection cannula (Phymep). Note that these empirically derived stereotaxic coordinates do not precisely match those given in the mouse brain atlas (Paxinos and Franklin, 2004), which we used as references for the injection-site images. To enable retrograde transport of the RB into the somas of midbrain DA neurons, we waited for an adequate time to perform the electrophysiology experiments, depending on the injection zone: 3 weeks after NAc injection and 2 weeks after Amg injection.

### Virus injection and optogenetics experiments

DAT^iCRE^ mice were anesthetized (Isoflurane 1%) and IC injection was performed into the VTA (coordinates bregma: + 3.1 mm; lateral: 0.5 mm; ventral: 4.5 mm) with 0.5 μl of an adeno-associated virus (AAV5.EF1α.DIO.CatCh.YFP 2.46e^12^ or 6.53e^14^ ng/μL, AAV5.EF1α.DIO.Jaws.eGFP 1.16e^14^ ng/μL diluted at 1:10, AAV5.EF1α.DIO.YFP 6.89e^14^ or 9.10e^13^ ng/μL). A double-floxed inverse open reading frame (DIO) allowed restraining the expression of CatCh (Ca^2+^ translocating channelrhodopsin) or Jaws (red-shifted cruxhalorhodopsin) to VTA DA neurons. Optical fibers (200 μm core, NA = 0.39, Thor Labs) coupled to a ferule (1.25 mm) were implanted bilaterally in the different target sites of the VTA (coordinates for BLA implantation: bregma: - 1.6 mm; lateral: 3.18 mm; ventral: 4.5 mm) (coordinates for NAc LatSh implantation: bregma: + 1.5 mm; lateral: 1.55 mm; ventral: 3.95 mm), and fixed to the skull with dental cement (SuperBond, Sun medical). An ultra-high-power LED coupled to a patch cord (500 μm core, NA = 0.5, Prizmatix) was used for optical stimulation (output intensity of 10 mW). Optical stimulation was delivered at a frequency of 10 Hz, 5 ms-pulse at 470 nm (Prizmatix LED) for activating stimulation, and with a continuous stimulation at 520 nm (Prizmatix LED) for inhibiting stimulation.

Optogenetic experiment performed on WT mice were conducted in the same manner as described above. Viruses were bilaterally injected into the VTA at the same coordinates as previously.

The first experiments were conducted on the animals at least 4 weeks after virus injection, to allow the construct to expressed by target cells. The optical stimulation cable was plugged onto the ferule during all experimental sessions to prepare the animals and control for latent experimental effects.

To perform non-specific optogenetic stimulation of different subnuclei of the amygdala (Amg), either AAV2-CAG-Jaws-GFP (1.45 e-12 ng/μL) or AAV2-7m8-CAG-GFP (5.70 e-12 ng/μL) were injected bilaterally into the VTA of distinct groups of wild-type (WT) mice as described in Methods. Bilateral optical fibers were implanted in those mice either in the basolateral amygdala (BLA, bregma: −1.6 mm; lateral: 3.18 mm; ventral: 4.5 mm) or in the central amygdala (CeA, bregma: −0.78 mm; lateral: 2.3 mm; ventral: 4.8 mm).

### Intracranial infusion

Bilateral guide cannulae (Bilaney) were implanted in the VTA (bregma: + 3.1 mm; lateral: 0.5 mm; ventral: 4.3 mm) of mice under anaesthesia (Isoflurane 3% induction and 1.5% maintenance) in order to allow local infusion of drugs prior to the EOM experiment. Before each experiment session, a double injection cannulae (4.5 mm length, 1 mm interval) was inserted into the implanted bilateral guide cannulae (length under pedestal 4.0 mm), 0.5 mm beyond the tip of the guide cannula. The injection cannula was connected to a multi-syringe pump (Univentor) that allowed saline or nicotine injection over 1 min (100 ng in 100 nl/injection).

### *In vivo* electrophysiology

Mice were deeply anaesthetized with chloral hydrate (8%), 400 mg/kg I.P., supplemented as required to maintain optimal anesthesia throughout the experiment. The scalp was opened and a hole was drilled in the skull above the location of the VTA. Intravenous administration of drugs was carried out through a catheter into the saphenous vein of the animal. Extracellular recording electrodes were constructed from 1.5 mm O.D. / 1.17 mm I.D. borosilicate glass tubing (Harvard Apparatus) using a vertical electrode puller (Narishige). The tip was broken straight and clean under microscopic control to obtain a diameter of about 1 μm. The electrodes were filled with a 0.5% NaCl solution containing 1.5% of neurobiotin^®^ tracer (VECTOR laboratories) yielding impedances of 6-9 MΩ. Electrical signals were amplified by a high-impedance amplifier (Axon Instruments) and monitored audibly through an audio monitor (A.M. Systems Inc.). The signal was digitized, sampled at 25 kHz, and recorded on a computer using Spike2 software (Cambridge Electronic Design) for later analysis. The electrophysiological activity was sampled in the central region of the VTA (coordinates: between 3.1 to 4 mm posterior to bregma, 0.3 to 0.7 mm lateral to midline, and 4 to 4.8 mm below brain surface). Individual electrode tracks were separated from one another by at least 0.1 mm in the horizontal plane. Spontaneously active DA neurons were identified based on previously established electrophysiological criteria (Grace and Bunney, 1984b; 1984a; Ungless and Grace, 2012).

After recording, nicotine-responsive cells were labelled by electroporation of their membrane. To do so, successive currents squares were applied until the membrane breaks in order to fill their soma with neurobiotin contained into the glass pipet (Pinault 1996). To be able to establish correspondence between neurons responses and their localization in the VTA, we labeled one type of response per mouse: solely activated neurons or solely inhibited neurons, with a limited number of cells per brain (1 to 4 neurons maximum, 2 by hemisphere), always with the same concern of localization of neurons in the VTA.

### *Ex vivo* patch-clamp recordings

For a functional verification of CatCh or Jaws expression, the same virus as described above was injected into 7-9 week-old male DAT^iCRE^ mice. For the characterization of NAc-projecting and Amg-projecting neurons, green retrobead tracers (Lumafluor) were injected into 7-9 weeks old male WT mice. After 4 weeks (for DAT-Cre) or 2 weeks (for WT), mice were deeply anesthetized by an intraperitoneal injection of a mix of ketamine (150 mg/kg Imalgene® 1000, Merial) and xylazine (60 mg/kg, Rompun® 2%, Bayer). Coronal midbrain sections (250 μm) were sliced with a Compresstome (VF-200, Precisionary Instruments) after intracardial perfusion of cold (4°C) sucrose-based artificial cerebrospinal fluid (SB-aCSF) containing (in mM): 125 NaCl, 2.5 KCl, 1.25 NaH_2_PO_4_, 5.9 MgCl_2_, 26 NaHCO_3_, 25 sucrose, 2.5 glucose, 1 kynurenate (pH 7.2, 325 mOsm). After 10 to 60 minutes at 35°C for recovery, slices were transferred into oxygenated aCSF containing (in mM): 125 NaCl, 2.5 KCl, 1.25 NaH_2_PO_4_, 2 CaCl_2_, 1 MgCl_2_, 26 NaHCO_3_, 15 sucrose, 10 glucose (pH 7.2, 325 mOsm) at room temperature for the rest of the day, and individually transferred to a recording chamber continuously perfused at 2 mL/minute with oxygenated aCSF. Patch pipettes (4-8 MΩ) were pulled from thin wall borosilicate glass (G150TF-3, Warner Instruments) with a micropipette puller (P-87, Sutter Instruments, Novato, CA) and filled with a potassium gluconate-based intracellular solution containing either (in mM): 116 K-gluconate, 20 HEPES, 0.5 EGTA, 6 KCl, 2 NaCl, 4 ATP, 0.3 GTP, and biocytin 2 mg/mL (pH adjusted to 7.2). Neurons were visualized using an upright microscope coupled with a Dodt contrast lens, and illuminated with a white light source (Scientifica). A 460 nm LED (pE-2, Cooled) was used for visualizing GFP, YFP- or RB positive cells (using a bandpass filter cube, AHF). Whole-cell recordings were performed using a patch-clamp amplifier (Axoclamp 200B, Molecular Devices) connected to a Digidata (1550 LowNoise acquisition system, Molecular Devices). Signals were low-pass filtered (Bessel, 2 kHz) and collected at 10 kHz using the data acquisition software pClamp 10.5 (Molecular Devices). Optical stimulation was applied through the microscope with two LEDs (460 nm and 525 nm, pE-2, CoolLED). To characterize CatCh expression, a 1 s continuous photostimulation was used to evoke currents in voltage-clamp mode (−60 mV), and a 10 Hz - 5 ms/pulse photostimulation was used to drive neuronal firing in current-clamp mode. Regarding Jaws expression, continuous photostimulation (20 sec) was used in current-clamp (−60 mV). To record nicotinic currents from RB+ DA neurons of the VTA, local puffs (500 ms) of nicotine tartrate (100 μM in aCSF) were applied with a glass pipette (2-3 μm diameter) positioned 20 to 30 μm away from the soma and connected to a picospritzer (World Precision Instruments, adjusted to ~2 psi). All electrophysiological recordings were extracted using Clampfit (Molecular Devices) and analyzed with R.

### Immunostaining

After euthanasia, brains were rapidly removed and fixed in 4% paraformaldehyde. After a period of at least three days of fixation at 4°C, serial 60-μm sections were cut from the midbrain with a vibratome. Immunostaining experiments were performed as follows: free-floating VTA brain sections were incubated for 1 hour at 4°C in a blocking solution of phosphate-buffered saline (PBS) containing 3% bovine serum albumin (BSA, Sigma; A4503) (vol/vol) and 0.2% Triton X-100 (vol/vol), and then incubated overnight at 4 °C with a mouse anti-tyrosine hydroxylase antibody (anti-TH, Sigma, T1299) and a chicken anti-eYFP antibody (Life technologies Molecular Probes, A-6455), both at 1:500 dilution, in PBS containing 1.5% BSA and 0.2% Triton X-100. The following day, sections were rinsed with PBS, and then incubated for 3 hours at 22-25 °C with Cy3-conjugated anti-mouse and Alexa488-conjugated anti-chicken secondary antibodies (Jackson ImmunoResearch, 715-165-150 and 711-225-152) at 1:500 and 1:1000 dilution in a solution of 1.5% BSA in PBS, respectively. After three rinses in PBS, slices were wet-mounted using Prolong Gold Antifade Reagent (Invitrogen, P36930). Microscopy was carried out with a fluorescent microscope, and images captured using a camera and analyzed with ImageJ.

In the case of electrophysiological recordings, the recorded neurons were identified by immunohisto-fluorescence as described above, with the addition of 1:200 AMCA-conjugated streptavidin (Jackson ImmunoResearch) in the solution. Immunoreactivity for both TH and neurobiotin (NB) allowed us to confirm the neurochemical phenotype of DA neurons in the VTA (TH+ NB+).

In the case of optogenetic experiments on DAT^iCRE^ mice, identification of the transfected neurons by immunohistofluorescence was performed as described above, with the addition of chicken-anti-eYFP primary antibody (1:500, ab13970, Abcam) in the solution. A goat-anti-chicken AlexaFluor 488 (1:500, Life Technologies) was then used as secondary antibody. Immunoreactivity for TH, YFP and neurobiotin/biocytin allowed us to confirm the neurochemical phenotype of DA neurons in the VTA (TH+ NB+) and the transfection success (YFP+).

### Images acquisition

All immunohistochemical slices were imaged by acquisition on a Leica DMR epi-fluorescent microscope, under identical conditions of magnification, illumination and exposure (using photometrics coolsnap camera). Images were captured in gray level using MetaView software (Universal Imaging Corporation, Ropper Scientific, France) and colored post-acquisition on ImageJ software.

### Behavioral data acquisition

All behavioral tests were conducted during the light period of the animal cycle (between 1:00 and 7:00PM). Experimental data were acquired using Anymaze software, coupled with 2D video camera.

#### Elevated O-maze (EOM) test

The EOM apparatus consists of two open (stressful) and two enclosed (protecting) elevated arms that together form a zero or circle (diameter of 50 cm, height of 58 cm, 10 cm-wide circular platform). Time spent in exploring enclosed versus open arms indicates then the anxiety level of the animal.

The first EOM experiment assess the effect of acute Nic injection (0.5mg/kg) on WT animals. The test lasts 10 minutes. Animals are injected 1-min prior the test and then put in the EOM for 9 min.

For intra-cranial infusion of Nic in the second experiment of EOM, animals received Nic (100ng/infusion) over 1-min prior the test. It lasted for 9 min as described above.

Finally, optogenetic experiments in EOM lasted for 15 minutes, with alternance of 5min-period of stimulation and non-stimulation (OFF-ON-OFF). When nicotine was injected in mice, the EOM test lasted for 9 min with the same protocol as described above.

#### *Online place preference* (OPP) *test*

The OPP protocols were performed in a Y-maze apparatus (Imetronic, Pessac, France), using only two arms of the Y-maze as two distinct compartments (the third arm was closed by a door and not available to the animal). The chamber in between is an equilateral triangle (side of 11 cm) used as a neutral compartment, where the animal was never photo-stimulated. Each arm of the maze measured 25 cm × 12 cm. The first arm displayed black and white stripes with smooth walls and floor, whereas the other arm displayed uniform-gray rough walls and floor. Choices of the compartment where the animals will be stimulated were counterbalanced across animals in the same test and YFP-control groups.

The OPP test consisted of a 20 minute-session where animals can freely navigate between the compartments but were photo-stimulated only in one of the two compartments.

Implanted bilateral fibers were connected with a bilateral fiber (diameter of 400 μm, NA = 0.39, Thorlabs) attached to a rotor connecting the 470 nm- or 520 nm-LED (Prizmatix) with a fiber of diameter 500 μm and NA = 0.5 (Thorlabs). LED output was controlled using a Master-8 pulse stimulator (A.M.P.I., Jerus) which delivered a discontinuous stimulation of 5-ms light flashes at 10 Hz frequency and 470 nm wavelength (for CatCh experiments), or a continuous stimulation at 520 nm (for Jaws experiments). Naive mice were connected and placed at the center of the neutral compartment before starting the recording.

#### Open field (OF) paradigm

The OF is a square enclosure of 50 cm × 50 cm where animals can move freely. Animal displacements were quantified by comparing the time spent in the center versus the periphery of the square. When nicotine was injected to WT mice in the OF test (IP injection of nicotine tartrate at 0.5 mg/kg, 0.1 mL/10 g, 1 minute prior to the test), animals were placed in the center of the OF for a 9-minute test duration, freely moving inside the enclosure. Regarding the optogenetic experiments conducted in the OF, animals were placed in the maze for 15 minutes, while alternating between OFF, ON and OFF optical stimulations of 5-minute periods.

### Data Analyses

#### Measurement of neuronal activity

Timestamps of action potentials were extracted in Spike 2 and analyzed using R, a language and environment for statistical computing (Team, 2005, http://www.r-project.org). Spontaneous activity of DA cell firing *in vivo* was analyzed with respect to the average firing frequency (in Hz) and the percentage of spikes-within-burst (%SWB = number of spikes within burst divided by total number of spikes in a given window). Neuronal basal activity was defined on at least five-minute recording.

#### Classification analysis and machine learning

To determine whether the spontaneous activity of in vivo-labeled, VTA DA neurons could predict their nicotine-evoked responses (activation or inhibition), we analyzed 8 variables that characterize the firing patterns: the mean firing frequency, the standard deviation and coefficient of variation of the firing frequency estimated on sliding windows, the percentage of spikes within bursts, the bursting frequency, the mean number of spikes in a burst, the fraction of time spent in bursting activity, and the burst length. Firing frequency and burst activity are quantified as described in the materials and methods section. Principal component analysis (PCA) and hierarchical clustering on principle components (HCPC) were conducted using R package FactoMineR. PCA and clustering distinguished 4 clusters that did not match with the nicotine-evoked responses. We then used the R package RandomForest to test for a prediction of nicotine-evoked responses from spontaneous cell activity. We determined that 5000 was the optimal number of trees in the forest and that 2 was the best number of variables randomly sampled as candidates at each split.

#### Quantification of nicotine responses

Firing frequency was quantified on overlapping 60-second windows shifted by 15 seconds time steps. For each neuron, firing frequency was rescaled as a percentage of its baseline value averaged during 3 minutes before nicotine injection. The responses to nicotine were thus presented as a percentage of variation from baseline (mean ± S.E.M.). The effect of nicotine was assessed by comparison of the maximum of firing frequency variation induced by nicotine and saline injection. For activated (respectively inhibited) neurons, the maximal (respectively minimal) value of the firing frequency was measured within the response period (3 minutes) that followed nicotine or saline injection. The results were presented as mean ± S.E.M. of the difference of maximum variation after nicotine or saline.

#### Subpopulations of VTA DA neurons in response to nicotine

Subpopulations of DA neurons were automatically classified using variation of firing frequency and the following routine: considering the maximal variation from a short baseline (3 minutes before injection), within 3 minutes after injection, neurons displaying an increase in firing frequency were defined as ‘‘activated’’, and neurons displaying a decrease in firing frequency were defined as ‘‘inhibited’’ (up or below zero in the cumulative distribution of responses to nicotine). In β2 ^−/−^ mice, VTA DA neurons did not show a clear change of their firing rate after nicotine injection (i.e., less than 5% of variation from baseline), and therefore were considered as a unique population to represent the mean response variation in Figure 3B.

#### Juxtacellular labeling

A total number of 245 neurons have been recorded and labeled for Figure 1. Those 245 neurons were used in Figure 1B-D. Among them, 101 neurons were shown in Figure 2B and E, containing 49 neurons labeled in NAc-RB injected mice and 52 in Amg-RB injected mice. The locations of the labeled neurons were manually placed on sections of the Paxinos atlas georeferenced in a 2D grid using adobe illustrator rules. The medio-lateral and dorso-ventral coordinates of the location of each neuron were extracted from the grid pattern and the antero-posterior coordinates were estimated from the section of the Paxinos atlas on which the neurons were placed. These three coordinates were used to make density histograms of location for nicotine-activated and nicotine-inhibited DA neurons or NAc-projecting and Amg-projecting DA neurons.

#### Behavioral experiments

The raw data for behavioral experiments were acquired as the time spent by animals in the different zones of the environments. The animals were detected in their body center by a USB camera connected to Anymaze software.

### Statistical analyses

All statistical analyses were done using the R software with home-made routines. Results are plotted as a mean ± S.E.M. The total number (n) of observations in each group and the statistics used are indicated in the figures directly or in the figure legends. Classically comparisons between means were performed using parametric tests as Student’s t-test, or two-way ANOVA for comparing two groups when parameters followed a normal distribution (Shapiro-Wilk normality test p > 0.05), or Wilcoxon non-parametric test as when the distribution was skewed. Bonferroni *post-hoc* analysis was applied, when necessary, to compare means. P > 0.05 was considered not to be statistically significant. Statistical significance was set at p < 0.05 (*), p < 0.01 (**), or p < 0.001 (***).

Figure 1: Kolmogorov-Smirnov test was used to compare the responses of VTA DA neurons to saline or nicotine injection. Wilcoxon tests were used to demonstrate a significant increase or decrease of firing frequency induced by nicotine injection compared to saline injection (B). Wilcoxon test was used to compared coordinates of nicotine-inhibited and nicotine-activated recorded neurons (C).

Figure 3: For behavior (A-C), over time effect of nicotine or saline injection (IP and IC) on the time spent by the mice in the open arms of the O-maze was first tested with one-way repeated measures ANOVA for each group of mice (shown in Figure S3.1B-D-G). Two-way repeated measures ANOVA (time/treatment or time/genotype) were used to compare the difference between the groups. In case of significant interaction effect between factors, Wilcoxon or Student’s t-test with Bonferroni corrections were used for intra- and inter-group *post-hoc* analysis (as indicated in the figure). For electrophysiology (B), Kolmogorov-Smirnov test was used to compare responses to nicotine of DA neurons in β2^−/−^ mice and β2^−/−^Vec mice. Wilcoxon tests with Bonferroni corrections are used to demonstrate a significant increase or decrease of firing frequency induced by IV nicotine injection in β2Vec mice compared to saline and nicotine injections in β2^−/−^ mice.

Figure 4: For O-maze experiments (A-C), effect of light was first tested with one-way repeated measures ANOVA for each group of mice (shown in Figure S4.3A-B-C). Two-way repeated measures ANOVA (epoch/opsin) were used to compare the difference between the groups. In case of significant interaction effect between factors, Wilcoxon or Student’s t-test with Bonferroni corrections were used for intra- and inter-group *post-hoc* analysis (as indicated in the figure). For O-maze experiment under nicotine (D), over time effect of nicotine injection on the time spent by the mice in the open arms of the O-maze was first tested with one-way repeated measures ANOVA for each group. Two-way repeated measures ANOVA (epoch/opsin) were used to compare the difference between the groups. In case of significant interaction effect between factor, Wilcoxon or Student’s t-test with Bonferroni corrections were used for intra- and inter- group *post-hoc* analysis (as indicated in the figure). For OPP experiments (E), preference score between group were compared with Student’s t-test.

Figure S2.1: Two-way repeated measures ANOVA (current, phenotype) is used to compare neuronal excitability (Fig. S2.1D). Wilcoxon test is used to compared coordinates of nicotine-inhibited and nicotine-activated recorded neurons (Fig. S2.1F).

Figure S2.2: Wilcoxon test is used to compare nicotine-evoked currents (Fig. S2.2F).

Figure S3: Two-way repeated measures ANOVA (time, treatment) is used to compare the distance traveled by mice in the open field (OF) after nicotine or saline injection (Fig. S3A). One-way repeated measures ANOVA is used to test the overtime effect of saline or nicotine intraperitoneal injection, or intracranial infusion, or the time spent by mice in the open arms of the elevated O-maze (EOM) (Fig. S3B, D, G). Two-way repeated measures ANOVA (time, genotype) is used to compare the time spent in the open arms of the EOM after nicotine injection between groups (Fig. S3F). In case of a significant interaction effect between factors, Wilcoxon or Student’s t-test with Bonferroni corrections is used for intra- and inter-group *post-hoc* analysis.

Figure S4.3: For anxiety, one-way repeated measures ANOVA is used to test the effect of light on the time spent in the open arms of the EOM (Fig. 4.3A-C). For locomotor activity, two-way repeated measures ANOVA (epoch, opsine) is used to compare the difference of light effect on the distance traveled by the mice between the groups (Fig. 4.3D-F). In case of a significant interaction effect between factors, Wilcoxon or Student’s t-test with Bonferroni corrections is used for intra- and inter-group *post-hoc* analysis (as indicated in the figure).

Figure S4.4: O-maze experiments (C-F) and locomotor activity (C-F) are analyzed as previously described for Fig.S4.3.

